# A Phosphorelay Circuit Drives Extracellular Alkalinization in Plant Receptor Kinase Signaling

**DOI:** 10.1101/2025.08.16.670655

**Authors:** Keran Zhai, Paul Derbyshire, Songyuan Zhang, Sera Choi, Limin Wang, Beibei Song, Toshinori Kinoshita, Jian-Min Zhou, Frank L. H. Menke, Kyle W. Bender, Cyril Zipfel

## Abstract

Extracellular alkalinization has long been recognized as a hallmark of plant cell-surface receptor activation, including during pattern-triggered immunity (PTI); yet the mechanisms driving elicitor-induced alkalinization and its role in immune signaling remain unclear. Here, we demonstrate that inhibition of autoinhibited H^+^-ATPases (AHAs) is required for elicitor-induced extracellular alkalinization. This alkalinization is essential for immune signaling mediated by diverse plasma membrane-localized receptor kinases (RKs) through modulation of ligand-receptor interactions. Notably, RKs transduce elicitor-triggered signaling via BOTRYTIS-INDUCED KINASE 1 (BIK1), which inhibits AHA activity by disrupting AHA-GENERAL REGULATORY FACTOR (GRF) interactions through a conserved phosphorylation event. Interestingly, this pathway is crucial for cell wall damage (CWD) responses involving the RK MALE DISCOVERER 1-INTERACTING RECEPTOR LIKE KINASE 2 (MIK2) and its ligand, SERINE RICH ENDOGENOUS PEPTIDE 18 (SCOOP18). Our findings reveal a conserved phospho-regulatory pathway that governs extracellular alkalinization to coordinate plant immune signaling, offering new insights into plant stress resilience.

## INTRODUCTION

All eukaryotes deploy cell-surface receptors to perceive external signals.^1,2^ In plants, plasma membrane (PM)-localized receptor kinases (RKs) play important roles in regulating growth, reproduction, and immunity.^2-4^ Among these, leucine-rich repeat receptor kinases (LRR-RKs) represent the largest subgroup.^2^ The extracellular LRR domain is responsible for recognizing specific ligands and is essential for the functional specificity of LRR-RKs.^5^ Two extensively characterized LRR-RKs in plants are the *Arabidopsis thaliana* (hereafter, Arabidopsis) FLAGELLIN SENSING 2 (FLS2) and ELONGATION FACTOR-TU RECEPTOR (EFR), which detect bacterial peptide epitopes flg22 (from flagellin) and elf18 (from Elongation Factor Tu), respectively, triggering pattern-triggered immunity (PTI).^6,7^ In addition to microbial ligands, LRR-RKs recognize endogenous plant-derived damage-associated molecular patterns (DAMPs) and immunomodulatory peptides, which function as phytocytokines.^8,9^ For instance, PLANT ENDOGENOUS PEPTIDE 1 (PEP1) RECEPTOR 1 (PEPR1) and PEPR2 detect PEP peptides to amplify immune responses,^10,11^ while MALE DISCOVERER 1-INTERACTING RECEPTOR-LIKE KINASE 2 (MIK2) recognizes SERINE-RICH ENDOGENOUS PEPTIDEs (SCOOPs) from both plants and microbes, contributing to immune regulation.^12-15^

Upon ligand perception, LRR-RKs commonly recruit the co-receptor BRASSINOSTEROID INSENSITIVE 1 (BRI1)-ASSOCIATED RECEPTOR KINASE 1 (BAK1), also known as SOMATIC EMBRYOGENESIS RECEPTOR-LIKE KINASE 3 (SERK3), along with other members of the SERK family.^5,16^ This co-receptor recruitment activates receptor complexes, leading to the phosphorylation of downstream receptor-like cytoplasmic kinases (RLCKs).^16,17^ Members of the RLCK-VII family, also known as AVRPPHB SUSCEPTIBLE 1 (PBS1)-LIKE KINASES (PBLs), play a critical role in mediating cytoplasmic signaling events.^18^ Among these, BOTRYTIS-INDUCED KINASE 1 (BIK1) and its homolog PBL1 are central signaling nodes downstream of diverse pattern recognition receptor (PRR) complexes, orchestrating PTI signaling.^17,18^

The activation of PTI triggers numerous cell signaling events, including ion flux changes, reactive oxygen species (ROS) production, mitogen-activated protein kinase (MAPK) activation, callose deposition, and extensive transcriptional reprogramming.^1,17,19^ Among these, extracellular alkalinization has been recognized as one of the earliest cellular responses, serving as a hallmark of PTI activation.^10,20-23^ However, the mechanisms by which plants induce extracellular alkalinization and how this alkalinization affects PTI activation remain elusive. Recent research has shed light on the role of PM-localized H^+^-ATPases, also known as autoinhibited H^+^-ATPase (AHA), in regulating extracellular pH.^24-29^ AHAs actively extrude protons from the cytosol, generating a transmembrane proton gradient and establishing a membrane potential.^30,31^ Increasing evidence indicates that phosphorylation of PM H⁺-ATPases plays a pivotal role in their regulation.^30,31^ The LRR-RK TRANSMEMBRANE KINASE 1 (TMK1) has been reported to phosphorylate the conserved penultimate Threonine residue (Thr^948^ in AHA1, Thr^947^ in AHA2), thereby activating AHA1 and AHA2, leading to apoplastic acidification and promoting auxin-mediated cell elongation.^25,26^ The plant secreted signaling peptide RAPID ALKALINIZATION FACTOR 1 (RALF1) induces extracellular alkalinization, which coincides with RALF1-triggered phosphorylation of AHA2 at Ser^899^, suggesting that AHA2 plays a critical role in RALF1-induced alkalinization.^29^ Phosphorylation at specific residues on AHAs modulates their binding affinity to 14-3-3 proteins, also known as GENERAL REGULATORY FACTORs (GRFs).^30,31^ These proteins stabilize the activated state of the AHAs, thereby regulating pH homeostasis.^30,31^ Several ligand-receptor pairs with pH-dependent characteristics have been identified in plants, with various biological functions.^24,32^ Notably, recent studies have characterized two pH-sensitive perception systems in roots: the ROOT MERISTEM GROWTH FACTOR 1 (RGF1) and its receptor (RGFR), as well as the PEP1-PEPR1 pair,^33^ highlighting the crucial regulatory role of pH in plant growth and immunity.

In this study, to comprehensively characterize the immune elicitor-triggered extracellular alkalinization, we selected flg22, elf18, and SCOOP18 as representative ligands for mechanistic analyses. We demonstrate that extracellular alkalinization is important for further immune signaling triggered by diverse elicitors, potentially by modulating ligand-receptor binding. Our findings reveal a previously uncharacterized pathway that mediates elicitor-triggered extracellular alkalinization, involving receptor/co-receptor complexes, BIK1, GRFs, and AHAs. Specifically, in the SCOOP18-MIK2 pathway, SCOOP18 induces MIK2-dependent phosphorylation of BIK1 at Tyr^250^, which is crucial for BIK1-mediated phosphorylation of GRFs at Thr^211^. This phosphorylation disrupts the interaction between GRFs and AHAs, thereby inhibiting AHA activity and leading to extracellular alkalinization. This pathway is also activated by other ligand-receptor pairs, such as flg22-FLS2 and elf18-EFR, and likely operates for additional alkalinization-inducing elicitors in plants. Notably, the phosphorylation site on GRFs is conserved across kingdoms, suggesting a common phospho-switch mechanism modulating pH changes during development and immunity. Importantly, this pathway is activated in response to cell wall damage (CWD), a common stress for plants, and is essential for CWD responses, highlighting its critical physiological role in plant defense and offering a potential strategy to enhance plant resilience to stresses.

## RESULTS

### Multiple immune elicitors alkalinize extracellular pH

To systematically investigate extracellular alkalinization triggered by immune elicitors, we selected flg22 and elf18 as representative non-self signals, and SCOOP18 as a representative host-derived altered self signal for further analysis.^6,7,13,14,34^ The kinetics of net H^+^ flux across the plasma membrane (PM) in single mesophyll cells, induced by these elicitors, were monitored using a non-invasive microelectrode (NMT)^35^. flg22, elf18, and SCOOP18 rapidly induced H^+^ influx in a manner dependent on their cognate receptors, FLS2, EFR and MIK2, respectively (Figures 1A-C). These experiments provided higher temporal and spatial (at PM) resolution than previous observations of flg22/elf18-induced alkalinization in cell suspension cultures^21,23^, suggesting that the extracellular alkalinization results from an increased net cellular H⁺ influx. Consistently, imaging of apoplastic pH using the membrane-impermeable, ratiometric, pH-sensitive fluorescent dye 8-hydroxypyrene-1,3,6-trisulfonic acid trisodium salt (HPTS)^36^ showed a significant increase in extracellular pH upon elicitor treatment (Figures 1D,E). This pH increase was also abolished in the corresponding receptor mutants (Figures 1F-K), indicating that extracellular alkalinization requires receptor-mediated signaling.

**Figure 1.**
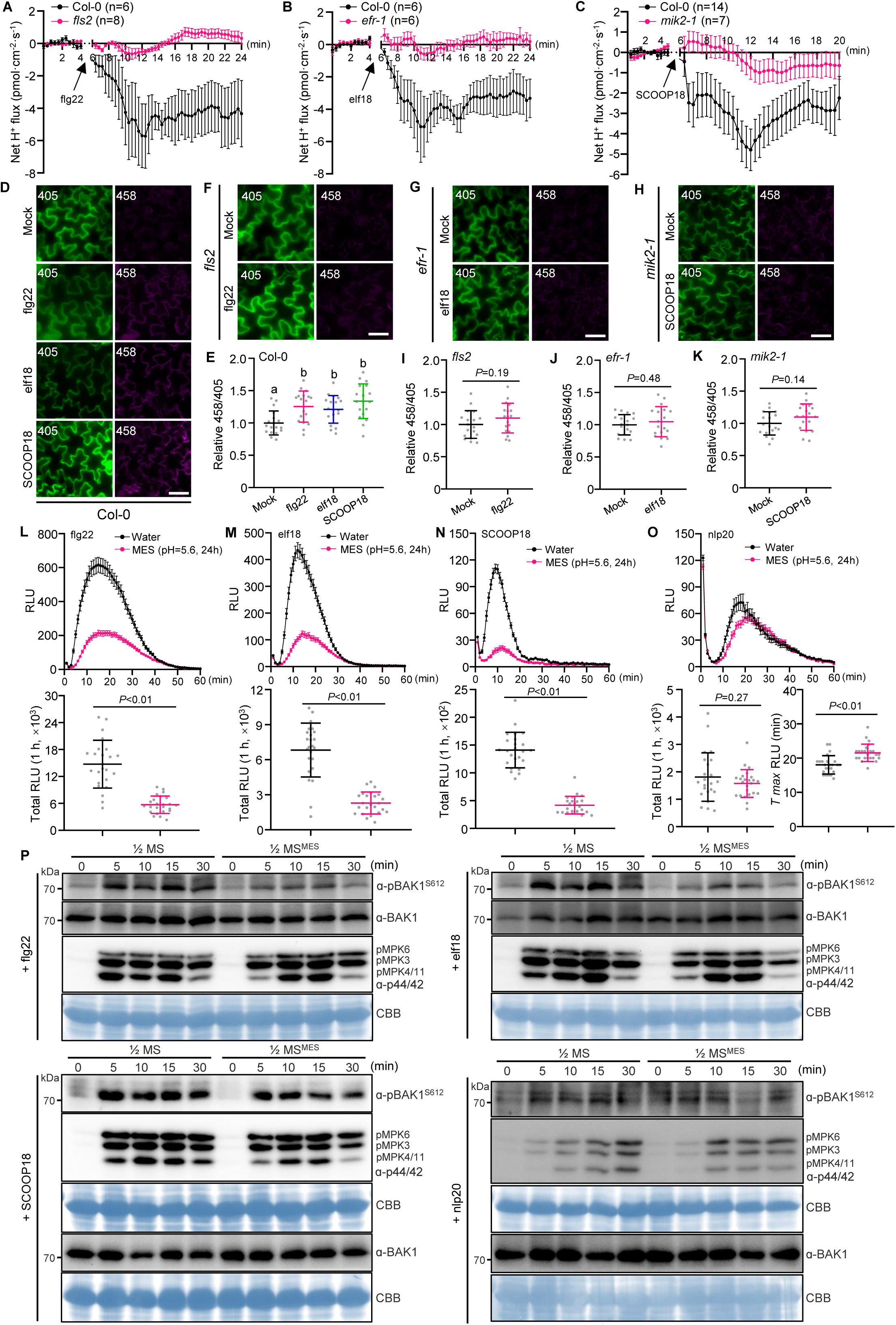
Elicitor-triggered extracellular alkalinization is essential for PTI signaling. (A-C) Net H^+^ flux kinetics upon flg22, elf18 and SCOOP18 treatment determined with NMT. Four-week-old mesophyll cells treated with flg22 (A), elf18 (B) or SCOOP18 (C). The data are shown as mean ± SE. (D and F-H) Representative confocal images of the HPTS-stained cotyledons of Col-0 (D), *fls2* (F), *efr-1* (G) or *mik2-1* (H) seedlings 1 h after treatment with flg22 (D, F), elf18 (D, G) or SCOOP18 (D, H). Scale bar, 50 μm. (E and I-K) Quantification of the mean 458 nm/405 nm ratios in elicitor-treated cells compared to the mock-treated cells is shown for (D), (F), (G) and (H), respectively. Two different forms of HPTS (protonated and deprotonated) were detected in two independent channels with excitation wavelengths of 405□nm and 458□nm, respectively. n□=□18 (6 independent seedling, 3 regions for each). (L-O) ROS production after treatment with different elicitors. Leaf discs of Col-0 were pretreated with or without 2 mM MES (pH 5.6) for 24 h, followed by treatment with flg22 (L), elf18 (M), SCOOP18 (N) or nlp20 (O). Values are means of RLU of independent leaf discs (upper panel), total RLU (down panel) or the time to maximum RLU (*T_max_* RLU) (lower right panel of O) over 1 h. The data are shown as mean ± SE (upper panel) or mean ± SD (down panel), n = 24. (P) BAK1 and MAP kinase phosphorylation in two-week-old seedlings of Col-0. Seedlings pretreated with or without 5 mM MES in ½ MS for 12 h, followed by treatment with flg22 (left, upper panel), SCOOP18 (left, down panel), elf18 (right, upper panel) or nlp20 (right, down panel). CBB was used as a loading control. 100 nM flg22, 100 nM elf18, 1 μM SCOOP18 or 1 μM nlp20 was used for the treatment. Different letters in (E) denote statistically significant differences according to one-way ANOVA followed by the Tukey’s test (*P* < 0.05). *P*-values indicate the data were analyzed by two-tailed Student’s *t*-test (I-N and lower left panel of O) or two-tailed Mann-Whitney test (lower right panel of O). The experiments were repeated at least twice (P) or three times (A-O) with similar results.

### Extracellular alkalinization is crucial for PTI signaling

To evaluate the effect of apoplastic pH on immune signaling, we manipulated extracellular pH by buffering with 2-(N-Morpholino) ethanesulfonic acid-KOH (MES-KOH, pH 5.6), a common buffer used in plant growth media, and assessed elicitor-triggered ROS production. Similar to what observed with PEP1, which has been reported to activate signaling in a pH-dependent manner,^33,37^ ROS production triggered by flg22 and SCOOP18 was markedly reduced under MES buffering conditions (Figures S1A-C), while ROS production induced by elf18 was significantly delayed (Figure S1D). In contrast, nlp20 (a 20-amino-acid peptide derived from bacterial, oomycete and fungal NECROSIS AND ETHYLENE-INDUCING PEPTIDE 1-LIKE PROTEINS), which does not induce alkalinization,^38^ showed no difference in ROS production (Figure S1E), indicating that neither the ROS production machinery nor the luminol pH sensitivity was affected by the pH manipulation. Notably, prolonged incubation in MES solutions resulted in a more pronounced inhibition of ROS production triggered by flg22, elf18, SCOOP18 and PEP1 (Figures 1L-N and S1F), while nlp20-triggered ROS production was only delayed, likely due to indirect effects of extended buffering (Figure 1O), suggesting that pH changes are critical for ROS production triggered by alkalinization-inducing elicitors.

Moreover, compared to normal conditions, the phosphorylation of BAK1^S612^, a marker for receptor activation,^39^ induced by flg22, elf18, and SCOOP18, but not by nlp20, was significantly reduced in MES-buffered conditions (Figure 1P). This reduction in phosphorylation was not due to impaired BAK1 accumulation (Figure 1P). While there were no significant changes with MPK3/6 activation between normal and MES-buffered conditions, the activation of MPK4/11 was reduced at the 5 min (Figure 1P), consistent with the previous report that BAK1^S612^ phosphorylation is crucial for MPK4/11 activation^39^.

We further investigated the apoplastic alkaline pH effect on flg22-, elf18-, and SCOOP18-triggered immune activation using different pH conditions. Treatment with a pH 8.0 solution did not induce ROS production on its own (Figure S1G), and a short incubation (∼1 min) at pH 8.0 did not affect ROS production triggered by flg22 (Figure S1H), indicating that the ROS measurement system is not impacted by this pH level. After pre-incubating leaves at different pH for 15 or 30 min, we found that, similar to what observed with PEP1 (Figure S1I), flg22- and SCOOP18-induced ROS production was significantly enhanced at pH 8.0 (Figures S1J,K). Although elf18 did not induce increased ROS production at pH 8.0, it occurred much faster (Figure S1L), suggesting different optimal pH ranges or sensitivities for different ligand-receptor pairs. This increased ROS production was not due to the addition of K^+^ (Figure S1M). Furthermore, pre-incubating plants at pH 8.0 resulted in enhanced BAK1 phosphorylation in response to flg22, elf18, and SCOOP18, whereas incubation at pH 4.1 led to a reduction in this phosphorylation (Figures S1N-P). Altogether, these results suggest that elicitor-triggered extracellular alkalinization modulates PTI signaling.

### Autoinhibited H^+^-ATPases (AHAs) are required for PTI signaling

The plant autoinhibited H^+^-ATPase family functions in mediating H^+^ efflux from the cytosol to the extracellular space,^31^ and has been implicated in auxin-induced acidification^25,26^. To explore the potential role of AHA inhibition in immune elicitor-triggered extracellular alkalinization, we selected AHA1, AHA2 and AHA11 for further analysis due to their similar expression patterns with FLS2, EFR and MIK2^7,13,40,41^. We first analyzed ROS production triggered by elicitors in an *aha1* T-DNA insertion null mutant (Figure S2A), since *aha1 aha2* double mutant is embryo lethal^42^. Interestingly, ROS production induced by flg22, elf18 and SCOOP18 was slightly but significantly reduced in *aha1* (Figures S2B-D), implying AHA1 is genetically involved in PTI signaling, potentially with redundant roles shared by AHA2 and AHA11.

To overcome this potential functional redundancy, we used virus-induced gene silencing (VIGS) to target *AHA2*, *AHA2/11*, or *GFP* as a control, in the *aha1* background. *AHA2* and *AHA11* transcripts were specifically and significantly downregulated in the *aha1/AHA-VIGS* plants (Figure S2E). Notably, *aha1/AHA2-VIGS* plants exhibited reduced growth, while *aha1/AHA2/AHA11-VIGS* plants displayed dwarfism compared to *aha1/GFP-VIGS* control (Figure S2F), highlighting the functional redundancy of AHA1, AHA2 and AHA11. flg22, elf18 and SCOOP18-induced ROS production was further reduced in *aha1/AHA2-VIGS* plants, with an even more pronounced reduction in *aha1/AHA2/AHA11-VIGS* plants (Figures 2A-C). Additionally, flg22- and elf18-triggered ROS production was largely reduced while SCOOP18-triggered ROS production was significantly delayed in the o*st2-1D* mutant,^43^ which harbours a constitutively activated AHA1 (Figures 2D-F). These findings suggest that AHAs are involved in PTI signaling, likely through their proton pump activity.

**Figure 2.**
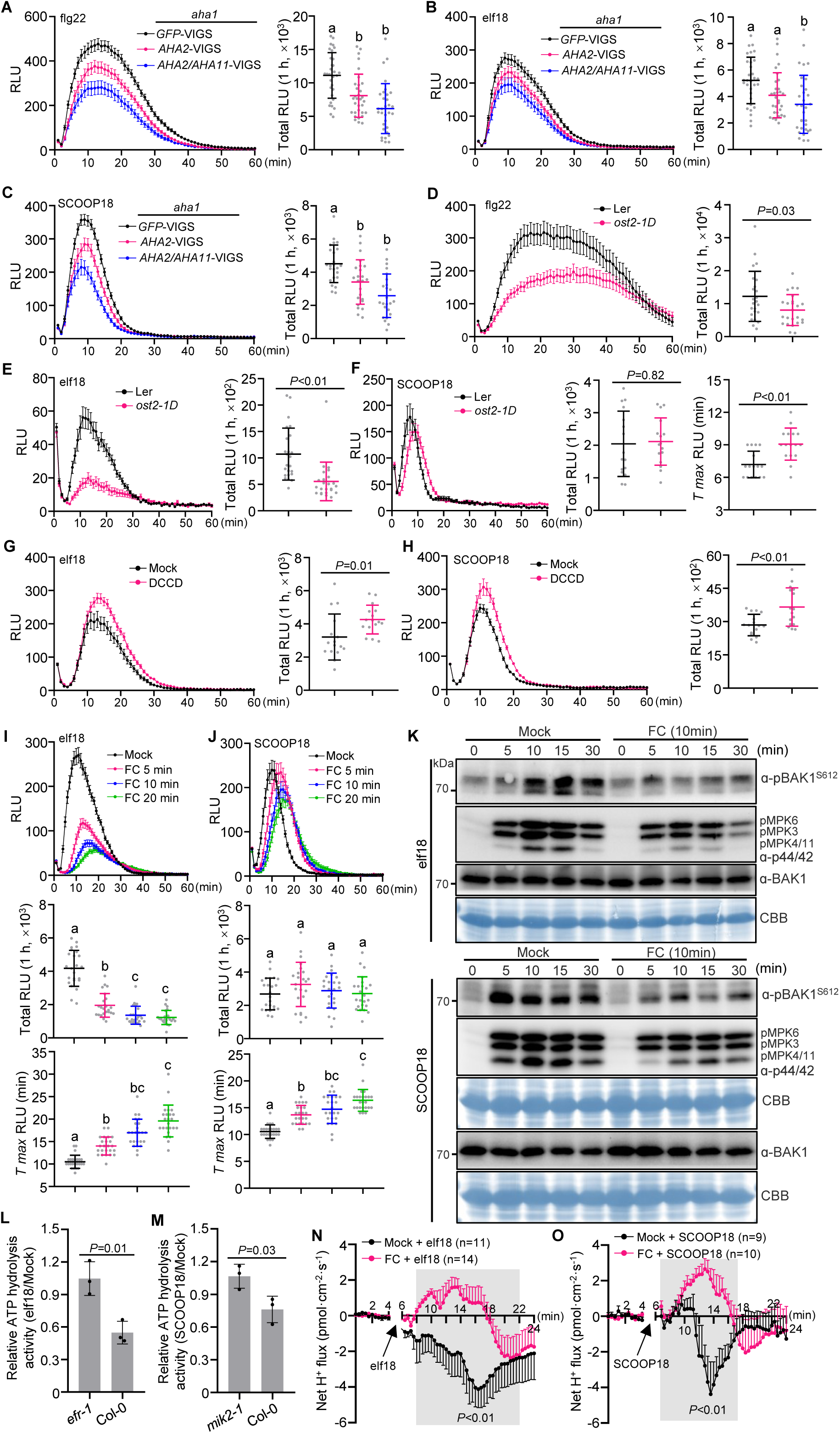
AHAs are required for elicitor-triggered H^+^ influx and PTI signaling. (A-C) ROS production after treatment with different elicitors. Leaf discs of *GFP*-VIGS, *AHA*2-VIGS and *AHA2/AHA11*-VIGS in *aha1* background were treated with flg22 (A), elf18 (B) or SCOOP18 (C). Values are means of RLU of independent leaf discs (left panel) or total RLU (right panel) over 1 h. The data are shown as mean ± SE (left panel) or mean ± SD (right panel), n = 32 (A and B), n = 24 (C). (D-F) ROS production after treatment with different elicitors. Leaf discs of L*er* and *ost2-1D* were treated with flg22 (D), elf18 (E) or SCOOP18 (F). Values are means of RLU of independent leaf discs (left panel), total RLU (right panels of D and E; middle panel of F) or the time to maximum RLU (*T_max_* RLU) (right panel of F) over 1 h. The data are shown as mean ± SE (left panel) or mean ± SD (total RLU and *T_max_* RLU), n = 24 (D and E), n = 16 (F). (G and H) ROS production after treatment with elf18 and SCOOP18. Leaf discs of Col-0 pretreated with 10 μM DCCD or mock for 2 h, and then treated with elf18 (G) or SCOOP18 (H). Values are means of RLU of independent leaf discs (left panel) or total RLU (right panel) over 1 h. The data are shown as mean ± SE (left panel) or mean ± SD (right panel), n = 16. (I and J) ROS production after treatment with elf18 and SCOOP18. Leaf discs of Col-0 pretreated with 5 μM FC for the indicated time points, and then treated with elf18 (I) or SCOOP18 (J). Values are means of RLU of independent leaf discs (upper panel), total RLU (middle panel) or *T_max_* RLU (down panel) over 1 h. The data are shown as mean ± SE (upper panel) or mean ± SD (middle and down panels), n = 24. (K) BAK1 and MPK phosphorylation in two-week-old seedlings of Col-0. Seedlings pretreated with 5 μM FC or mock for 10 min, followed by treatment with elf18 (upper panel), or SCOOP18 (down panel). CBB was used as a loading control. (L and M) Quantification of ATP hydrolysis activity in Col-0, *efr-1* and *mik2-1*. Ten-day-old seedlings treated with elf18 (L) or SCOOP18 (M) for 5 min and then collected for ATP hydrolysis activity measurement. (N and O) Net H^+^ flux kinetics upon elf18 or SCOOP18 treatment determined with NMT. Four-week-old mesophyll cells were co-treated with elf18 and 5 μM FC (N) or SCOOP18 and 5 μM FC (O), with elf18 and DMSO (N) or SCOOP18 and DMSO (O) used as controls. The data are shown as mean ± SE.100 nM flg22, 100 nM elf18 or 1 μM SCOOP18 was used for the treatment. Different letters in (A-C and the middle panel of J) denote statistically significant differences according to one-way ANOVA followed by the Tukey’s test (*P* < 0.05). Different letters in (I and the down panel of J) denote statistically significant differences according to Kruskal-Wallis test followed by Dunn’s test (*P* < 0.05). *P*-values indicate the data were analyzed by two-tailed Mann-Whitney test (E and the right panel of F), two-tailed Student’s *t*-test (D, G, H, L, M and the middle panel of F) or two-way ANOVA (shaded areas of N and O). The experiments were repeated at least twice (K, N and O) or three times (A-J, L and M) with similar results.

Unexpectedly, both loss- and gain-of-function of AHA1 resulted in reduced immune signaling (Figures 2A-F), implying that the appropriate dynamic regulation of AHA activity and associated pH transients are critical for immune signaling. To test this hypothesis, the proton pump inhibitor N, N’-dicyclohexylcarbodiimide (DCCD) and the pump activator fusicoccin (FC) were employed to introduce transient pH changes.^33^ Immune responses were assessed using elf18 as a non-self signal, which induced a significantly reduced ROS production in *ost2-1D* (Figure 2E), and SCOOP18 as a self-derived signal. Interestingly, DCCD treatment enhanced elf18- and SCOOP18-induced ROS production (Figures 2G,H), while FC treatment inhibited elf18-triggered ROS and delayed SCOOP18-triggered ROS (Figures 2I,J), indicating AHA-mediated H^+^ efflux interferes with elicitor-triggered ROS production. Consistently, elf18- and SCOOP18-triggered BAK1 and MAPK (in particular MPK4/11) phosphorylation were reduced following FC treatment and increased following DCCD treatment (Figures 2K and S2G). These results support the notion that PTI signaling relies on finely tuned, AHA-mediated transient pH changes.

### AHAs are essential for elicitor-triggered H^+^ influx and resulting extracellular alkalinization

To determine whether elicitor treatment affects AHA activity, we performed an ATP-hydrolysis assay measuring the hydrolytic release of inorganic phosphate from ATP, representing AHA activity^44^. Interestingly, we detected decreased ATP hydrolysis activity after elf18 and SCOOP18 treatment in wild-type Col-0 (Figures 2L and 2M). This inhibition was absent in *efr-1* and *mik2-1* (Figures 2L,M), indicating that the elicitor-triggered inhibition of AHA activity is dependent on receptor-mediated signaling. Importantly, constitutive activation of AHA by FC treatment prevented the early-stage H^+^ influx typically induced by elf18 and SCOOP18, resulting in a delayed and weakened H^+^ influx at later time-points (Figures 2N,O), which could be attributed to other H^+^ transporters or unknown compensatory mechanisms. In addition, elf18- and SCOOP18-triggered extracellular alkalinization was abolished upon FC co-treatment (Figures S2H,I). Notably, the H⁺ influx triggered by elf18 and SCOOP18 was also reduced in *AHA1/AHA2/AHA11*-VIGS plants (Figures S2J-M), consistent with their impaired immune signaling activation (Figures 2B and 2C). Altogether, these findings suggest that transient inhibition of AHA activity is required for elicitor-induced net H^+^ influx and subsequent extracellular alkalinization.

### MIK2 inhibits AHA activity through BIK1

Phosphorylation is required for extracellular alkalinization following elicitor perception, as evidenced by complete inhibition of elicitor-triggered extracellular alkalinization in the presence of the phosphorylation inhibitor K252a.^20^ To investigate the mechanism underlying AHA inhibition after elicitor treatment, we focused on MIK2 due to its stronger interaction with AHA1 compared to EFR and FLS2 (Figures S3A-C). Although MIK2 consititutively forms a complex with AHA1, AHA2, and AHA11 *in vitro* and *in vivo* (Figures S3D-J), it does not directly phosphorylate the cytosolic regions of AHA1, AHA2, or AHA11 (Figures S3K-M).

Perception of SCOOP peptides triggers the association of MIK2 and its co-receptors BAK1/BAK1-LIKE 1 (BKK1) and relays the signaling through the receptor-like cytoplasmic kinases BIK1 and PBL1.^12-15,46-48^ Consistent with this, SCOOP18-induced ROS production was reduced in *bik1-2* and further diminished in *bik1-10 pbl1-10* mutants generated by CRISPR (Figures 3A and S4A). Additionally, it was abolished in *bak1-5 bkk1-1* (Figure 3A), indicating that BAK1/BKK1 and BIK1/PBL1 are essential for SCOOP18-triggered signaling. Surprisingly, SCOOP18-triggered inhibition of ATPase activity however remained unaffected in *bak1-5 bkk1-1* (Figure 3B) but was nearly abolished in *bik1-2* and *bik1-10 pbl1-10* (Figure 3C), suggesting that BIK1/PBL1, but not BAK1/BKK1, are required for AHA inhibition. Collectively, these findings indicate that MIK2 interacts with AHAs and inhibits their activity through BIK1.

**Figure 3.**
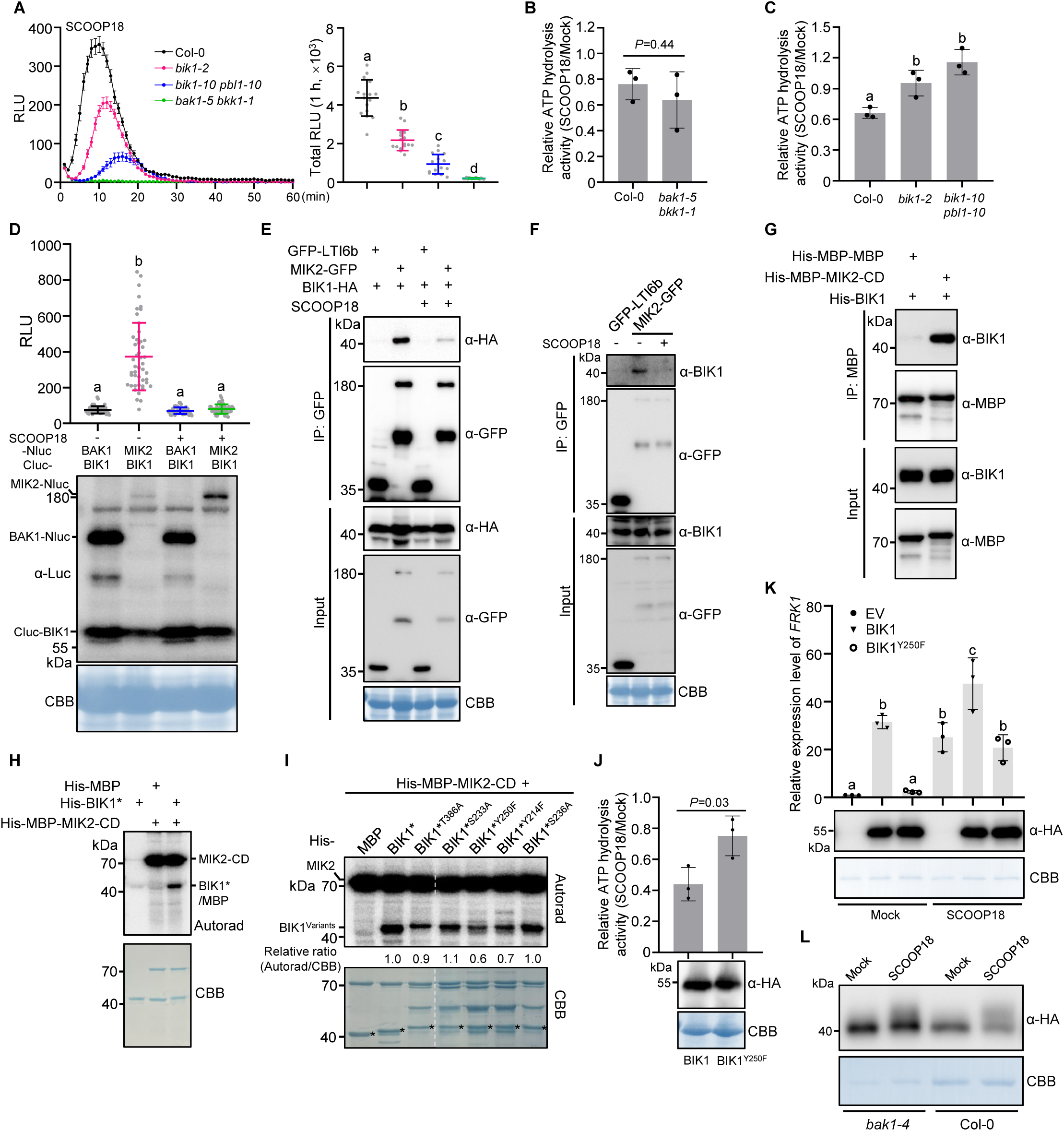
MIK2 interacts with and phosphorylates BIK1 to regulate AHA activity. (A) ROS production after treatment with SCOOP18. Leaf discs of Col-0, *bik1-2*, *bik1-10 pbl1-10* and *bak1-5 bkk1-1* were treated with SCOOP18. Values are means of RLU of independent leaf discs (left panel) or total RLU (right panel) over 1 h. The data are shown as mean ± SE (left panel) or mean ± SD (right panel), n = 16. (B and C) Quantification of ATP hydrolysis activity in Col-0, *bak1-5 bkk1-1, bik1-2* and *bik1-10 pbl1-10*. Ten-day-old seedlings treated with SCOOP18 or mock for 5 min and then collected for ATP hydrolysis activity measurement. (D) MIK2-BIK1 interaction in *N. benthamiana*. SLC assays were conducted with the indicated constructs in the presence or absence of 1 μM SCOOP18 for 10 min. Protein expression was detected by western blot (down panel). BAK1-BIK1 was used as a negative control. Values are means of RLU of independent leaf discs ± SD, n = 48. (E) Co-immunoprecipitation of MIK2-GFP and BIK1-HA transiently expressed in *N*. *benthamiana* leaves treated with SCOOP18 or mock for 10 min. The plasma membrane marker GFP-LTI6b was used as a negative control. (F) Co-immunoprecipitation of MIK2 and BIK1 in stable transgenic plants *mik2-1/35S::MIK2-GFP*. Seedlings were treated with SCOOP18 or mock for 10 min. Endogenous BIK1 protein was detected in immunoblots using α-BIK1 antibody. The GFP-LTI6b transgenic seedlings were used as a negative control. (G) MBP pulldown of His-BIK1 by MBP-MIK2-CD. The His-MBP-MBP was used as a negative control. (H) The MIK2-CD phosphorylates BIK1* (BIK1^K105A/K106A^) *in vitro*. Autoradiogram of *in vitro* kinase assay using His-MBP-MIK2-CD with His-BIK1*. His-MBP was used as a negative control. (I) *In vitro* phosphorylation assays for analyzing different residues in BIK1*. MIK2-CD was incubated with different BIK1* variants. His-MBP was used as a negative control. Relative ratio of ^32^P incorporation was quantified using ImageJ and is indicated under the lanes. The asterisks denote the corresponding proteins. White dashed lines denote cropping positions on the same membrane. (J) Quantification of ATP hydrolysis activity. BIK1 or BIK1^Y250F^ plasmids were transfected into *bik1-2* protoplasts, then treated with mock or SCOOP18 for 5 min, and collected for ATP hydrolysis activity measurement. Protein expression was detected by western blot (down panel). (K) Activation of *FRK1* by BIK1 variants. Empty vector (EV), BIK1, and BIK1^Y250F^ plasmids were transfected into *bik1-2* protoplasts, treated with mock or SCOOP18 for 1.5 h. Protein expression was detected by western blot (down panel). (L) SCOOP18-induced BIK1 mobility shift in protoplasts. Protoplasts from Col-0 or *bak1-4* were transfected with BIK1-2×HA and treated with SCOOP18 for 10 min. 1 μM SCOOP18 was used for the treatment (A-F and J-L). Different letters in (A, C, D and K) indicate statistically significant differences (*P* < 0.05), determined by one-way ANOVA with Tukey’s test (A, C and K) or Kruskal-Wallis test with Dunn’s test (D). *P*-values indicate the data were analyzed by two-tailed Student’s *t*-test (B and J). CBB indicates loading control (D-F and J-L) or loading of the protein (H and I). The experiments were repeated at least twice (H and I) or three times (A-G and J-L) with similar results.

### MIK2 interacts with and phosphorylates BIK1

The genetic independence of AHA inhibition from BAK1/BKK1 prompted us to test whether BIK1 directly interacts with MIK2. Split-luciferase complementation (SLC) and co-immunoprecipitation (Co-IP) assays in *Nicotiana benthamiana* revealed that MIK2 interacts with BIK1 (Figures 3D,E). This association was strongly reduced upon SCOOP18 treatment (Figures 3D,E), suggesting that BIK1 is released from the receptor complex upon SCOOP18 perception, similar to what was observed previously after flg22 perception by FLS2^49,50^. BIK1-MIK2 association and SCOOP18-induced dissociation were further confirmed in Arabidopsis *mik2-1/35S::MIK2-GFP* transgenic plants using anti-BIK1 antibody (Figure 3F). Additionally, an *in vitro* pull-down assay revealed that MIK2 directly interacts with BIK1 through its cytosolic kinase domain (Figure 3G).

We next tested whether MIK2 could directly phosphorylate BIK1 to transduce SCOOP18-triggered signaling. An *in vitro* radioactive kinase assay demonstrated direct phosphorylation of His-BIK1* (kinase-dead variant) by His-MBP-MIK2-CD (cytosolic domain) (Figure 3H). Further mass spectrometry analysis identified four phosphorylation sites on BIK1* (Y214, S233, Y250, and T386) detected in at least two independent replicates (Table S1). Among these, the phospho-ablative Y250F variant exhibited reduced phosphorylation by MIK2 in *in vitro* radioactive kinase assays, compared to the wild-type and other phospho-ablative variants (Figure 3I). As Y250 is required for BIK1 function in flg22-triggered signaling,^51^ we decided to prioritize its characterization. Similarly, overexpression of BIK1, but not BIK1^Y250F^, in *bik1-2* mesophyll protoplasts rescued SCOOP18-triggered inhibition of AHA activity and induction of the immune marker gene *FRK1* (Figures 3J,K), indicating Y250 is an essential phosphorylation site of BIK1 for SCOOP18-induced H^+^ influx and signaling activation. Moreover, SLC and Co-IP assays revealed that the kinase activity of MIK2 is not required for its interaction with BIK1, whereas it is indispensable for SCOOP18-induced dissociation (Figures S4B,C). Notably, SCOOP18-induced BIK1-MIK2 dissociation still occurred with BIK1^Y250F^ mutant (Figure S4D), suggesting that Y250 is not essential for BIK1 release from the MIK2 receptor complex upon SCOOP18 perception. It is plausible that other phosphorylation sites on BIK1, targeted by MIK2, also contribute to SCOOP18-triggered signaling.

Consistent with direct phosphorylation of BIK1 by MIK2, SCOOP18-induced BIK1 band shift (indicative of phosphorylation^50^) still occurred in *bak1-4* (Figure 3L), whereas BAK1-dependent BIK1 phosphorylation in response to flg22 treatment was abolished in *bak1-4*^50^ (Figure S4E). Considering that BAK1 forms a complex with MIK2 upon SCOOP perception,^12-15,46-48^ it is likely that BAK1 plays little direct role in BIK1 phosphorylation, which instead directly occurs through MIK2. It is also possible that other SERK family members have redundant functions with BAK1 for BIK1 phosphorylation. Altogether, these findings suggest that MIK2 interacts with and phosphorylates BIK1 to regulate SCOOP18-triggered extracellular alkalinization, while the canonical MIK2-BAK1 complex regulates other aspects of PTI signaling, such as ROS production.

### BIK1 promotes dissociation of GRF2 from AHA1 upon SCOOP18 perception

We next investigated how BIK1 interferes with AHA activity, focusing on AHA1 due to its strong interaction with BIK1 (Figure S4F). Although BIK1 directly interacts with AHA1 and this association is disrupted by SCOOP18 treatment (Figures S4G-I), no direct phosphorylation of AHA1 or AHA2 by BIK1 was detected *in vitro* (Figures S4J,K), suggesting that BIK1-mediated inhibition of AHAs occurs through a mechanism other than direct phosphorylation.

The interaction between 14-3-3s (GRFs) and AHAs is critical for activating AHAs.^30,31^ Given that BIK1 can directly interact with and inhibit AHA activity, we next sought to determine whether GRF-AHA interaction is affected by BIK1. GRF2 was selected for further analysis due to its specific enrichment in our AHA1-GFP immunoprecipitation coupled with mass spectrometry (IP-MS) dataset (Figure S5A and Table S2), and its role as a representative member of the non-□ group of 14-3-3 proteins, which are known to be more effective in stimulating H^+^-ATPase activity than □ group members (e.g., GRF9 and GRF10, as also observed in our IP-MS dataset)^52^. As expected, GRF2 interacted with AHA1 in the SLC assay (Figure 4A). Strikingly, this interaction was abolished when BIK1, but not BIK1* was co-expressed (Figure 4A), indicating that BIK1 inhibits GRF2-AHA1 interaction through its kinase activity, aligning with its physiological role. Additionally, the inhibition effect was compromised with BIK1^Y250F^ compared to BIK1 (Figure 4A), suggesting MIK2-mediated phosphorylation on BIK1 is important for inhibiting AHA activity. It is possible that the regulation mediated by Y250 might involve the control of BIK1 kinase activity. The effects of BIK1, BIK1* and BIK1^Y250F^ on GRF2-AHA1 interaction were further confirmed with Co-IP assays in Arabidopsis (Figure 4B). Notably, GRF2-AHA1 interaction was decreased upon SCOOP18 treatment in a MIK2-dependent manner in *N. benthamiana* (Figure S5B), reminiscent of BIK1-mediated inhibition of GRF2-AHA1 interaction. In line with this, Co-IP assays in Arabidopsis mesophyll protoplasts revealed that SCOOP18-triggered inhibition of GRF2-AHA1 interaction was largely dependent on BIK1 (Figure 4C), a finding further validated in *bik1-10 pbl1-10* (Figure 4D). Altogether, these results suggest that BIK1 inhibits AHA activity by interfering with AHA-GRF interaction.

**Figure 4.**
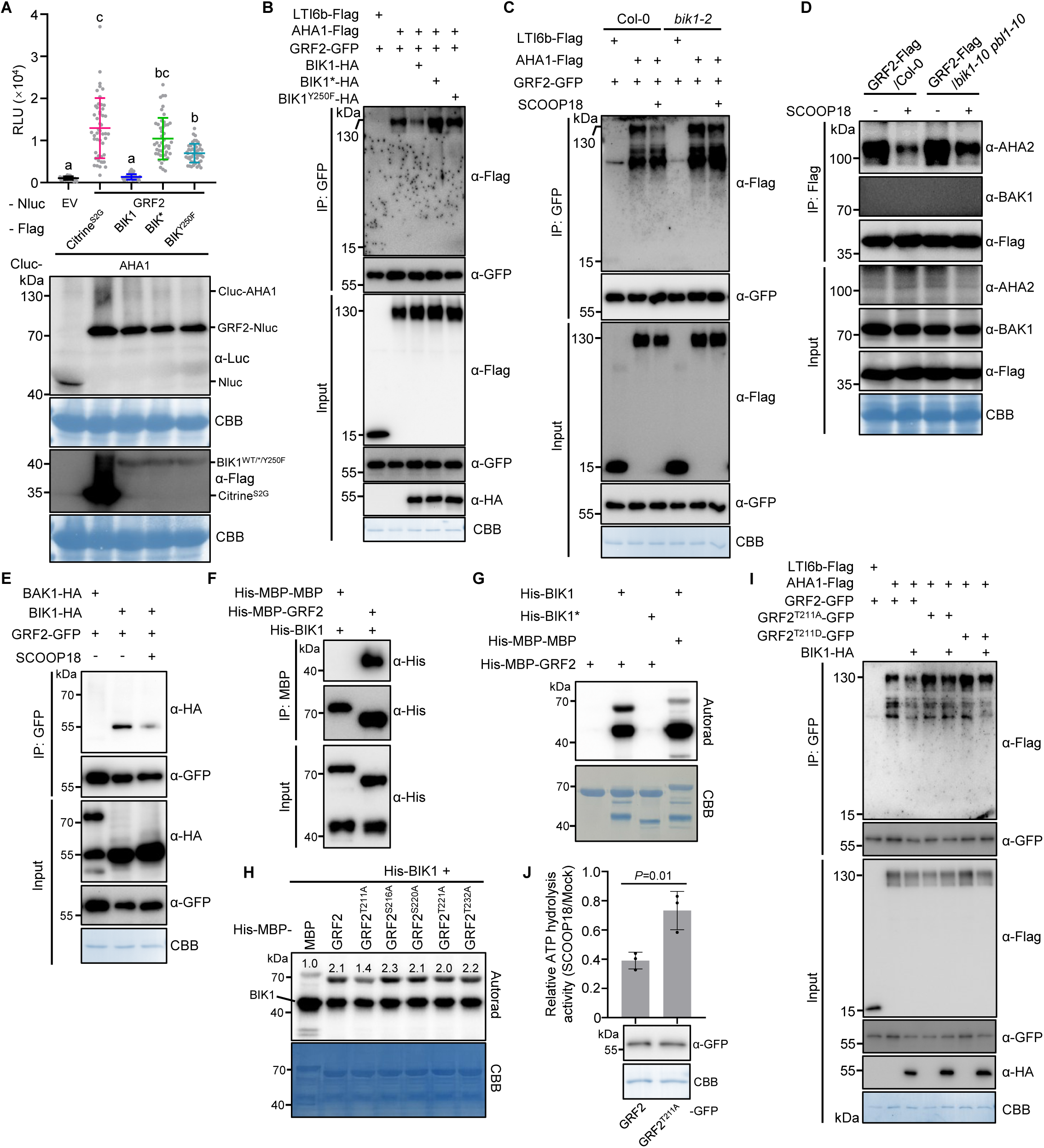
Phosphorylation of GRF2 by BIK1, leading to its dissociation from AHA1, thereby inhibiting AHA activity. (A) Effects of BIK1 variants on AHA1-GRF2 interaction in *N*. *benthamiana*. SLC assays were conducted with the indicated constructs. Protein expression was detected by western blot (down panel). The empty vector (Nluc) and Citrine^S2G^-Flag were used as negative controls. Values are means of RLU ± SD, n = 24 or 48 independent leaf discs. Different letters denote statistically significant differences according to Kruskal-Wallis test followed by Dunn’s test (*P* < 0.05). (B and C) Co-immunoprecipitation of AHA1 and GRF2 in protoplasts. Protoplasts from *bik1-2* were transfected with the indicated constructs (B). Note that 6×HA tagged BIK1^WT/*/Y250F^ were used. Protoplasts from Col-0 or *bik1-2* were transfected with the indicated constructs and then treated with 1 μM SCOOP18 or mock for 10 min (C). The LTI6b-Flag was used as a negative control. (D) Co-immunoprecipitation of AHA2 and GRF2-Flag in transgenic plants. Seedlings were treated with 1 μM SCOOP18 or mock for 10 min. Endogenous AHA2 protein was detected in immunoblots using α-AHA2 antibody. The endogenous BAK1 was used as a negative control. (E) Co-immunoprecipitation of BIK1 and GRF2 in protoplasts. Protoplasts from Col-0 were transfected with the indicated constructs and then treated with 1 μM SCOOP18 or mock for 10 min. The BAK1 was used as a negative control. (F) MBP pulldown of His-BIK1 by His-MBP-GRF2. The His-MBP-MBP was used as a negative control. (G) BIK1 phosphorylates GRF2 *in vitro*. Autoradiogram of *in vitro* kinase assay with the indicated protein combination. His-BIK1* and His-MBP-MBP were used as negative controls. (H) *In vitro* phosphorylation assays for analyzing different residues in GRF2. BIK1 was incubated with different GRF2 variants. His-MBP-MBP was used as a negative control. Relative ratio of ^32^P incorporation (Autorad/CBB) was quantified using ImageJ and is indicated above the bands. (I) Interaction analysis between BIK1 and different GRF2 variants using Co-IP assays. Protoplasts from Col-0 were transfected with the indicated constructs. The LTI6b-Flag was used as a negative control. (J) Quantification of ATP hydrolysis activity. GRF2 or GRF2^T211A^ plasmids were transfected into Col-0 protoplasts, then treated with mock or 1 μM SCOOP18 for 5 min, and collected for ATP hydrolysis activity measurement. Protein expression was detected by western blot (down panel). *P*-values indicate the data were analyzed by two-tailed Student’s *t*-test. CBB indicates loading control (A-E, I and J) or loading of the protein (G and H). The experiments were repeated at least twice (B and D) or 3 times (A, C and E-J) with similar results.

### BIK1 interacts with and phosphorylates GRF2 at Thr^211^

Given that BIK1 inhibits the GRF2-AHA1 interaction through its kinase activity but BIK1 cannot phosphorylate AHA1, we speculated that this inhibition might result from the phosphorylation of GRF2 by BIK1. To test this hypothesis, we first examined the interaction between BIK1 and GRF2. Co-IP assays in Arabidopsis revealed that BIK1 interacts with GRF2 (Figure 4E). Interestingly, this association was reduced upon SCOOP18 treatment (Figure 4E), suggesting that GRF2 dissociates from BIK1 upon SCOOP18 perception, which aligns with previous reports that interactions between BIK1 and its substrates are dynamically regulated during immune activation^53-56^. In addition, an *in vitro* pull-down assay revealed that BIK1 directly interacts with GRF2 (Figure 4F).

We next sought to determine whether GRF2 is a BIK1 substrate. An *in vitro* radioactive kinase assay revealed that BIK1 directly phosphorylates GRF2 (Figure 4G). Further mass spectrometry analysis identified five GRF2 residues (Thr^211^, Ser^216^, Ser^220^, Thr^221^ and Thr^232^) being trans-phosphorylated *in vitro* by BIK1, but not by BIK1* (Table S3). Notably, the Thr^211^-containing phospho-peptides exhibited the highest abundance and were consistently detected in all three replicates (Table S3). In line with this, the phosphorylation level of the phospho-ablative variant GRF2^T211A^ by BIK1 was reduced compared to wild type and the other four phospho-ablative variants in *in vitro* radioactive kinase assays (Figure 4H), suggesting that Thr^211^ is a major site for BIK1-mediated trans-phosphorylation on GRF2. Interestingly, Thr^211^ is not present within the Ser-X-X-Leu motif previously reported for several BIK1 substrates,^53,54,56-64^ suggesting that BIK1 might differentially phosphorylate distinct substrate motifs as is known for other protein kinases^65,66^. The mechanistic basis for such substrate switching remains an open question.

Notably, a recent quantitative phospho-proteomics study showed that flg22 treatment induces Thr^211^ phosphorylation of GRF2 in Col-0 plants,^67^ consistent with our observations and supporting the notion of elicitor-induced Thr^211^ phosphorylation *in vivo*.

### Phosphorylation of GRF2 at Thr^211^ is required for its complete dissociation from AHA1

To determine whether GRF2^Thr211^ phosphorylation is involved in BIK1-mediated inhibition of GRF2-AHA1 interaction, we compared the interactions of phospho-ablative (GRF2^T211A^) or phospho-mimetic (GRF2^T211D^) mutant variants with AHA1. Interestingly, we found that GRF2^T211A^ exhibited a stronger interaction with AHA1 compared to wild-type GRF2 in Arabidopsis protoplasts (Figure 4I), which was further corroborated in transgenic plants (Figure S5C). This enhanced interaction is likely due to the absence of phosphorylation at Thr^211^ of GRF2 by endogenous BIK1, in line with the increased interaction between GRF2 and AHA1 in *bik1-2* compared to Col-0 (Figure 4C). Notably, the inhibition of the AHA1-GRF2 interaction upon BIK1 overexpression was significantly reduced with GRF2^T211A^ (Figure 4I), suggesting that BIK1 inhibits this interaction mainly through Thr^211^. Surprisingly, GRF2^T211D^ showed a similar interaction with AHA1 as GRF2^T211A^ (Figures 4I and S5C), indicating this mutation may not accurately mimic the phosphorylated form of GRF2, in analogy to what has been reported for GRF6^68^. Furthermore, SCOOP18-triggered inhibition of AHA activity and production of ROS were significantly reduced when GRF2^T211A^ was expressed compared to wild-type GRF2 (Figures 4J and S5D), highlighting the physiological significance of Thr^211^ phosphorylation in the regulation of AHA activity and immune signaling.

To further investigate whether the residual inhibition of the AHA1-GRF2^T211A^ interaction by BIK1 depends on its kinase activity and potentially involves other phosphorylation sites, we co-expressed BIK1 or BIK1* with GRF2^T211A^, GRF2^T211A/S216A/S220A/T221A^ (GRF2^4A^), or GRF2^T211A/S216A/S220A/T221A/T232A^ (GRF2^5A^) and AHA1 in Arabidopsis protoplasts. We observed that, compared to BIK1*, BIK1 still inhibited the AHA1-GRF2^T211A^ interaction (Figure S5E). Similar inhibition was also observed with GRF2^4A^ or GRF2^5A^ (Figure S5F), indicating that the remaining inhibition of the AHA1-GRF2^T211A^ interaction by BIK1 is dependent on its kinase activity and independent of the other four identified phosphorylation sites. It is possible that other unknown components or compensatory mechanisms involved in the AHA1-GRF2 interaction are also regulated by BIK1 through its kinase activity. Taken together, the data support that phosphorylation of GRF2 at Thr^211^ by BIK1 contributes to its full dissociation from AHA1, thereby inhibiting AHA activity.

### GRF2 Thr^211^ is conserved across kingdoms

Notably, we found that Thr^211^ is conserved in other Arabidopsis GRF members, except for GRF13, and is substituted by Ser in GRF10 and GRF11 (Figure S6A). This conservation suggests a shared phosphorylation mechanism between BIK1 and GRFs. GRF10 was selected for testing this hypothesis due to its phylogenetic divergence from GRF2 and its enrichment in the AHA1-GFP IP-MS dataset (Figures S6A and S5A). *In vitro* radioactive kinase assays revealed that BIK1 directly phosphorylates GRF10 mainly in a Ser^209^-dependent manner (Figures S6B and S6C), where Ser^209^ in GRF10 corresponds to Thr^211^ in GRF2. Importantly, phosphorylation at Ser^209^ of GRF10 was also induced by flg22 treatment, as detected in a quantitative phospho-proteomics dataset from Col-0 plants,^67^ suggesting that BIK1-mediated phosphorylation at Thr^211^ on GRFs likely represents a conserved mechanism in immune signaling.

Strikingly, the Thr^211^ phosphorylation site and its flanking amino acid sequence in GRF2 are highly conserved among 14-3-3 proteins across kingdoms, including all plant species analyzed here, as well as in *Drosophila melanogaster*, *Danio rerio*, *Mus musculus* and *Homo sapiens* (Figure S6D). Notably, in human, the corresponding site in 14-3-3ε, Thr^208^, is phosphorylated by the acidophilic Polo like kinase 2/3 (PLK 2/3),^69^ although the functional implications remain unknown. Given this sequence conservation, it is plausible that other species might employ a similar phospho-switch regulatory mechanism to modulate pH changes during development and/or immunity.

### Inhibition of AHA activity by BIK1 is conserved in response to elicitors

To further investigate the conservation of BIK1/PBL1 in elicitor-triggered inhibition of AHA activity, ATP-hydrolysis assays were conducted. The results revealed that, like SCOOP18, flg22/elf18-triggered inhibition of AHA activity was abolished in *bik1-10 pbl1-10* (Figures S6E,F), suggesting a conserved function role for BIK1/PBL1 in this process. Surprisingly, unlike SCOOP18 (Figure 3B), flg22/elf18-triggered AHA inhibition was also abolished in *bak1-5 bkk1-1* (Figures S6G,H). This differential involvement of BAK1/BKK1 in flg22/elf18- and SCOOP18-regulated extracellular alkalinization was further confirmed by H^+^ flux measurements using NMT (Figure S6I). Combined with previous biochemical evidence showing that FLS2 and EFR are unable to phosphorylate BIK1 at Tyr^250^,^50,70^ while their co-receptor BAK1 can,^51^ we speculate that, although the BIK1/PBL1-GRF2-AHA1 module is conserved in its regulation by different elicitors, the upstream kinases involved in the phosphorylation relay may differ.

### pH changes impact PTI signaling by regulating ligand-receptor binding

To systemically understand how extracellular alkalinization contributes to PTI activation, we first tested whether it functions through the extracellular or intracellular domains of receptors using rapamycin (Rap) system^71^. In this system, the extracellular domain of the receptor is removed, and Rap-induced dimerization of the intracellular domains of isolated receptors (FLS2, EFR, MIK2) and the co-receptor BAK1 is sufficient to activate downstream signaling.^71,72^ Consistent with results in Arabidopsis, ROS production triggered by flg22, elf18, and SCOOP18 was significantly reduced under MES-buffered conditions in *N. benthamiana* (Figures 5A-C and S7A), whereas Rap-induced ROS production remained unaffected (Figures 5D-F and S7B). Additionally, FC treatment significantly inhibited flg22-, elf18-, and SCOOP18-triggered ROS production (Figures 5G-I and S7C), but did not inhibit Rap-induced ROS production (Figures 5J-L and S7D). Surprisingly, Rap-induced ROS production was even enhanced in the presence of EFR/BAK1 and MIK2-BAK1 (Figures 5K,L), likely due to intracellular pH changes. Furthermore, manipulation of apoplastic pH using alkaline solutions significantly enhanced flg22- and SCOOP18-triggered ROS production and accelerated elf18-triggered ROS production (Figures S7C and E-G), but had no effect on Rap-induced ROS production (Figures S7D and H-J). These observed effects are not due to alterations in the ROS production machinery or luminol pH sensitivity, as evidenced by the unchanged ROS production in the Rap system. These findings suggest that pH changes modulate FLS2-, EFR-, and MIK2-mediated signaling activation through their extracellular domains.

**Figure 5.**
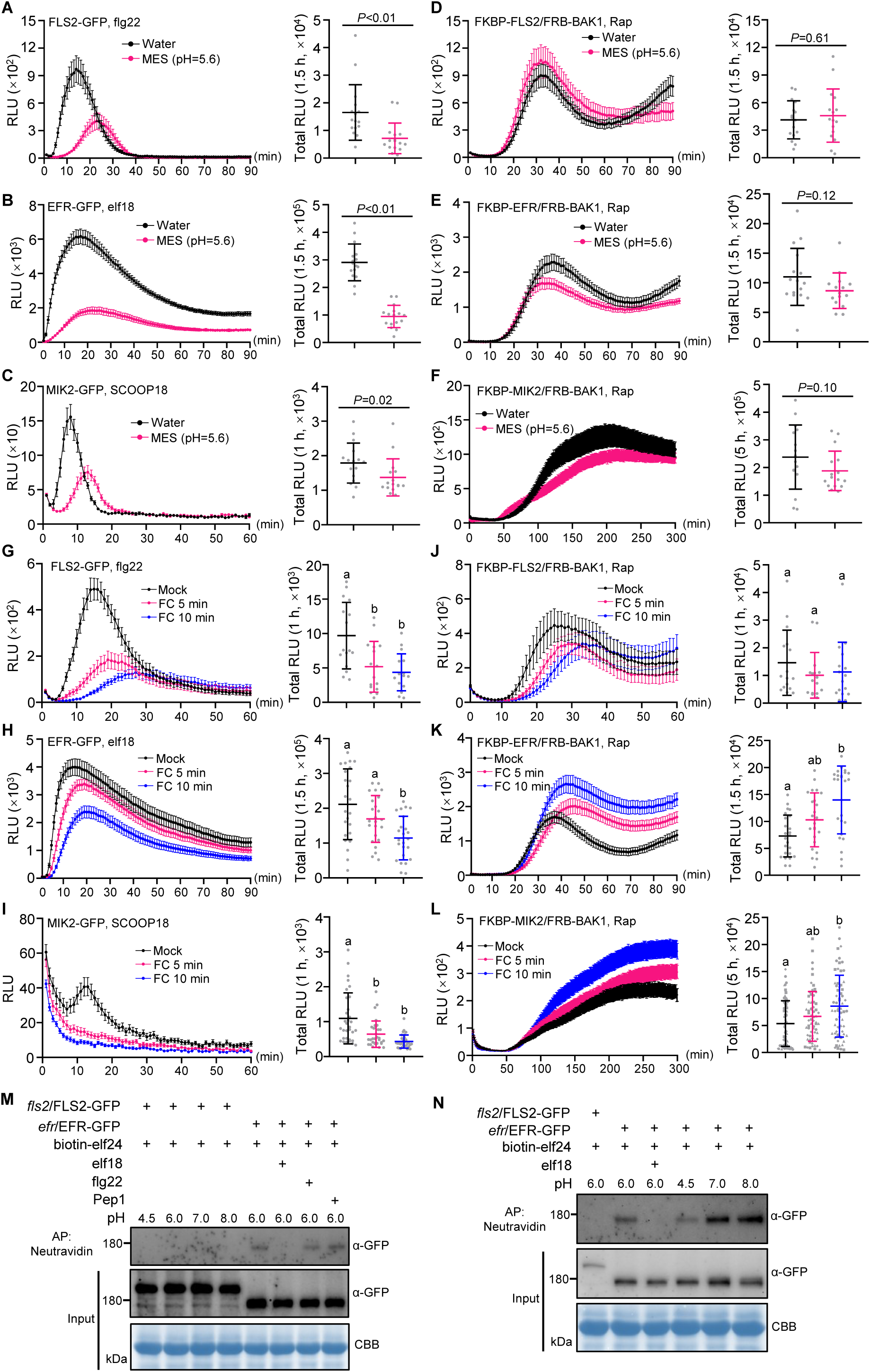
pH changes impact PTI activation via the receptor extracellular domains and affect ligand-receptor binding. (A-C) ROS production in *N. benthamiana* after treatment with different elicitors. Leaf discs of *N*. *benthamiana* leaves transiently expressing FLS2, EFR, or MIK2 were pretreated with or without 2 mM MES (pH 5.6) for 4 h, followed by treatment with flg22 (A), elf18 (B) or SCOOP18 (C). Values are means of RLU of independent leaf discs (left panel) or total RLU (right panel) over the indicated time. The data are shown as mean ± SE (left panel) or mean ± SD (right panel), n = 16. (D-F) ROS production in *N. benthamiana* induced by Rap. Leaf discs of *N*. *benthamiana* leaves transiently expressing FLS2/BAK1, EFR/BAK1, or MIK2/BAK1 were pretreated with or without 2 mM MES (pH 5.6) for 4 h, followed by treatment with Rap. Values are means of RLU of independent leaf discs (left panel) or total RLU (right panel) over the indicated time. The data are shown as mean ± SE (left panel) or mean ± SD (right panel), n = 16. (G-I) ROS production in *N. benthamiana* after treatment with different elicitors. Leaf discs of *N*. *benthamiana* leaves transiently expressing FLS2, EFR, or MIK2 were pretreated with FC for the indicated time, followed by treatment with flg22 (G), elf18 (H) or SCOOP18 (I). Values are means of RLU of independent leaf discs (left panel) or total RLU (right panel) over the indicated time. The data are shown as mean ± SE (left panel) or mean ± SD (right panel), n = 16 (G), 24 (H) or 32 (I). (J-L) ROS production in *N. benthamiana* after treatment with Rap. Leaf discs of *N*. *benthamiana* leaves transiently expressing intracellular domains of FLS2/BAK1, EFR/BAK1, or MIK2/BAK1 were pretreated with FC for the indicated time, followed by treatment with Rap. Values are means of RLU of independent leaf discs (left panel) or total RLU (right panel) over the indicated time. The data are shown as mean ± SE (left panel) or mean ± SD (right panel), n = 16 (J), 24 (K) or 64 (L). Four independent experiments represented by different dot shapes were merged into one graph (L). (M and N) *In vitro* binding assay of EFR-GFP with biotin-labelled elf24. Pull-down was performed using Neutravidin beads under the indicated pH conditions. Immunoblots show the amount of EFR-GFP or FLS2-GFP protein from Arabidopsis transgenic plants (input) and after affinity purification (AP). FLS2-GFP was used as a negative control. 1 μM biotin-elf24, 100 μM elf18, 100 μM flg22 or 100 μM Pep1 was used for the binding assays. 100 nM flg22, 100 nM elf18, 1 μM SCOOP18, 5 μM FC or 1 μM Rap was used for the treatment (A-L). Different letters in (G and H) denote statistically significant differences according to one-way ANOVA followed by the Tukey’s test (*P* < 0.05). Different letters in (I-L) denote statistically significant differences according to Kruskal-Wallis test followed by Dunn’s test (*P* < 0.05). *P*-values indicate the data were analyzed by two-tailed Mann-Whitney test (A, C and F) or two-tailed Student’s *t*-test (B, D and E). CBB indicates loading control (M and N). The experiments were repeated at least three times with similar results.

To test whether the ligand-receptor binding is affected by pH, like PEP1-PEPR1^33,37^, we performed binding assays using elf24 and EFR^73^. Biotinylated elf24 specifically bound to EFR-GFP, but not to FLS2-GFP, and this interaction was competitively inhibited by non-labeled elf18, but not by unrelated flg22 or PEP1 (Figure 5M). Interestingly, high pH (7.0 and 8.0) significantly promoted the binding of biotinylated elf24 to EFR, while low pH (4.5) reduced this interaction (Figure 5N). This suggests that elf24-EFR binding is pH-dependent, with alkalinization promoting binding and acidification inhibiting it. Combined with the biochemical and genetic data (Figures 5 and S7), we speculate that this pH preference may also apply to flg22-FLS2 and SCOOP18-MIK2 binding, which warrants further investigation. Together, these data suggest that extracellular alkalinization promotes PTI activation likely through enhancing ligand-receptor binding.

### Cell wall damage induced by cellulose biosynthesis inhibition activates MIK2-BIK1-AHA-mediated H^+^ influx

We recently reported that SCOOP18 is involved in cell wall integrity (CWI) signaling induced upon cellulose biosynthesis inhibition (CBI).^34^ To explore the biological relevance of SCOOP18-induced alkalinization in response to cell wall damage (CWD), we used isoxaben (ISX), which induces rapid clearing of cellulose synthase complexes from the PM, leading to CBI, mimicking CWD.^74,75^ Interestingly, ISX-induced responses, such as ROS production, lignification, and immune marker gene expression, were significantly reduced or completely abolished under MES-buffered conditions (Figures 6A-C), suggesting a direct involvement of pH changes in CBI-CWD responses. Notably, like SCOOP18, ISX treatment also induced H^+^ influx (Figure 6D), resulting in extracellular alkalinization (Figure 6E), reminiscent of *SCOOP18* induction after ISX treatment^34^. Consistently, ISX-induced extracellular alkalinization was significantly reduced in *mik2-1* (Figure 6E). Furthermore, ISX-induced H^+^ influx was significantly reduced in *mik2-1* and *bik1-2* (Figures 6D,F), and was abolished in *aha1/AHA2/11-VIGS* plants (Figure 6G), suggesting that the MIK2-BIK1-AHA signaling pathway also contributes to extracellular alkalinization triggered by CBI-CWD. Additionally, ISX treatment resulted in decreased AHA activity in wild-type Col-0, which was significantly reduced in *mik2-1* and *bik1-2* (Figures 6H,I), suggesting that CBI-CWD inhibits AHA activity through MIK2 and BIK1. Consistently, ISX-induced lignification and immune-marker gene expression were reduced in *bik1-2* (Figures 6J,K), and ISX-induced ROS production was abolished in *aha1/AHA2/11-VIGS* plants (Figure 6L), indicating BIK1 and AHA1 are genetically involved in CBI-CWD responses. Taken together, these findings, along with the previous report,^34^ demonstrate the biological significance of the SCOOP18-MIK2-BIK1-AHA cascade-mediated extracellular alkalinization in response to CWD, a common environmental stress in plants.

**Figure 6.**
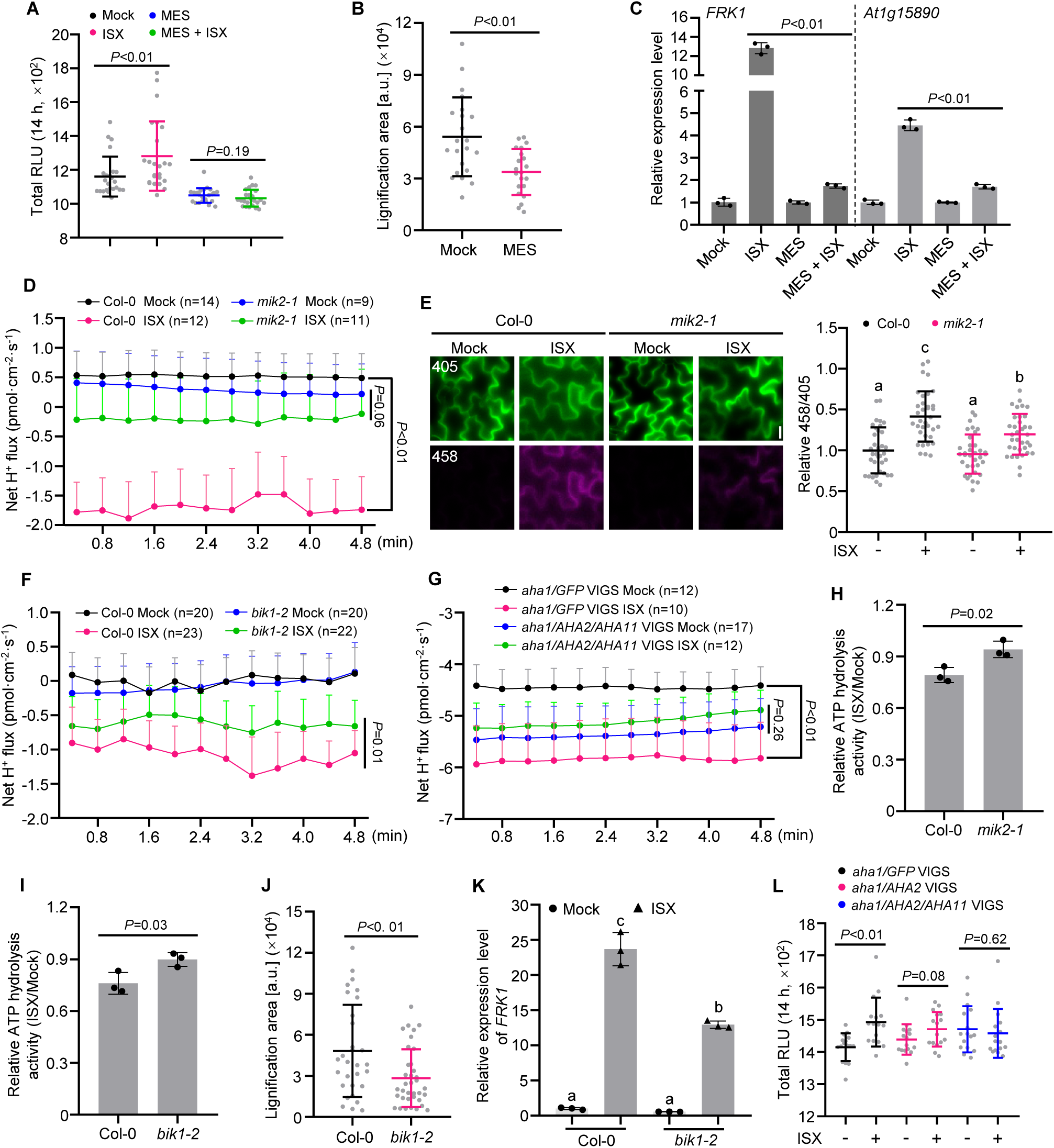
MIK2-BIK1-AHAs mediated H^+^ influx is required for CWD responses. (A) ISX-triggered ROS production. Leaf discs of Col-0 were treated with 1 μM ISX in water (mock) or MES buffer (pH 5.6). Values are means of total RLU over 14 h ± SD, n = 24. (B) Quantification of ISX-induced lignification. Root lignification in Col-0 seedlings after treatment with 0.6 μM ISX in water (mock) or MES buffer (pH 5.6) for 12 h was visualized by phloroglucinol staining. a.u., arbitrary units. Values are means of lignification areas ± SD, n = 22 (mock), n = 21 (MES). (C) RT-qPCR analysis of *FRK1* and *At1g15890* expression levels. Seedlings of Col-0 were treated with ISX in water (mock) or MES buffer (pH 5.6) for 12 h. *P*-values indicate the data were analyzed by two-tailed Student’s *t*-test. Data are mean ± SD (n = 3). (D) Net H^+^ flux kinetics after ISX treatment. Mesophyll cells from four-week-old Col-0 or *mik2-1* were treated with 1 μM ISX for approximately 20 h and H^+^ flux was measured with NMT. (E) Confocal images showing the apoplastic pH in cotyledon cells treated with ISX. Five-day-old seedlings treated with 0.6 μM ISX for 36 h and stained with HPTS. Changes in pH (imaging: left panel; quantification: right panel, mean ± SD, n = 35) were visualized with ratiometric values of fluorescent HPTS. (F) Net H^+^ flux kinetics after ISX treatment. Four-week-old mesophyll cells from Col-0 and *bik1-2* treated with 1 μM ISX for approximately 20 h and then measured with NMT. Data are mean ± SE. (G) Net H^+^ flux kinetics after ISX treatment. Four-week-old mesophyll cells of *GFP*-VIGS and *AHA2/AHA11*-VIGS in *aha1* background treated with 1 μM ISX for approximately 20 h and then measured with NMT. (H and I) Quantification of ATP hydrolysis activity in Col-0, *mik2-1* and *bik1-2*. Ten-day-old seedlings treated with 0.6 μM ISX or mock for 12 h and then collected for ATP hydrolysis activity measurement. (J) Quantification of ISX-induced lignification. Root lignification in Col-0 and *bik1-2* seedlings after treatment with 0.6 μM ISX for 12 h was visualized by phloroglucinol staining. Lignification area was quantified using ImageJ. a.u., arbitrary units. Values are means of lignification areas ± SD, (Col-0, n = 30), (*bik1-2*, n = 36). (K) RT-qPCR analysis of *FRK1* expression levels in Col-0 and *bik1-2*. Two-week-old seedlings treated with 0.6 μM ISX for 12 h. (L) ISX-triggered ROS production. Leaf discs of *GFP*-VIGS, *AHA2*-VIGS and *AHA2/AHA11*-VIGS in *aha1* background were treated with 1 μM ISX. Values are means of total RLU over 14 h ± SD, n = 16. *P*-values indicate the data were analyzed by two-tailed Mann-Whitney test (A and J), two-tailed Student’s *t*-test (B, C, H, I and L) or two-way ANOVA (D, F and G). Different letters in (E and K) denote statistically significant differences according to one-way ANOVA followed by the Tukey’s test (*P* < 0.05). The experiments were repeated at least twice (C and K) or three times (A, B, D-J and L) with similar results.

## DISCUSSION

Elicitor-triggered extracellular alkalinization in plants was first observed in 1991,^22^ and has since been widely recognized as a hallmark of plant defense responses.^10,20-22^ Despite this, two longstanding and fundamentally important questions remain unresolved: (1) How do plant cells induce extracellular alkalinization upon elicitor perception?, and (2) How does extracellular alkalinization influence PTI signaling? In this study, we aim to systematically investigate the underlying mechanisms, with a particular focus on how extracellular alkalinization is achieved, while also providing valuable insights into the regulation of PTI signaling mediated by extracellular alkalinization, using three representative elicitors as models, namely flg22, elf18 and SCOOP18. Our findings support a model (Figure 7) in which upon ligand perception, the receptor directly phosphorylates BIK1/PBL1 (e.g., for SCOOP18-MIK2) or does so indirectly via its co-receptor BAK1/BKK1 (e.g., for flg22-FLS2 and elf18-EFR). BIK1 subsequently phosphorylates GRFs, promoting their dissociation from AHAs, which inhibits AHA activity and triggers extracellular alkalinization. This alkalinization enhances ligand-receptor binding, thereby activating and/or amplifying downstream signaling. We have thus identified a phospho-switch pathway comprising the receptor/co-receptor-RLCK-GRF-AHA axis, which regulates extracellular alkalinization triggered by multiple elicitors. Given the conservation of these components^18,31,76^ and their phosphorylation sites across plant species (Figure S6D), this module could be widely utilized by plants to fine-tune pH changes in response to various elicitors under stress conditions. We further demonstrate that extracellular alkalinization is crucial for PTI signaling (Figures 1 and S1), potentially by enhancing the binding affinity between the receptor and its ligand (Figures 5M,N). Additionally, we provide evidence for the biological significance of the BIK1-GRFs-AHAs cascade-mediated extracellular alkalinization in response to CWD in the context of SCOOP18-MIK2 signaling (Figure 6), a common environmental stress in plants^77^. Interestingly, some studies suggest that pathogens can suppress plant immunity by lowering the environmental pH,^78,79^ indicating that apoplastic pH manipulation may serve as a battleground between plants and pathogens. Notably, dynamic pH changes are induced by various stresses, including drought and salt, as well as by developmental responses.^80,81^ Whether a similar regulatory mechanism is employed by plant to fine-tune pH changes during growth and abiotic stresses needs further investigation.

**Figure 7.**
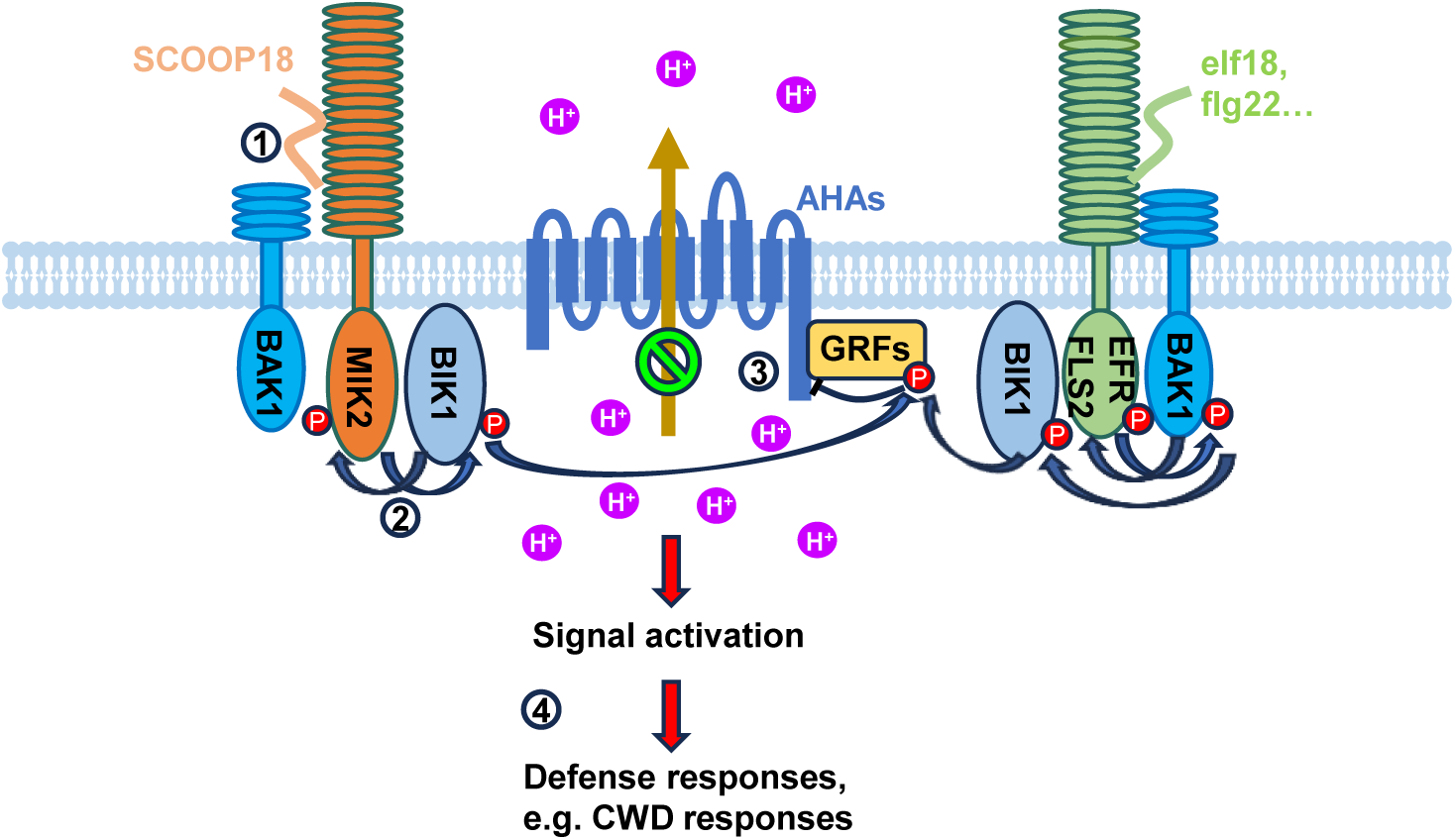
A proposed model for extracellular alkalinization triggered by plant cell-surface receptor kinases in defense activation. Upon exposure to biotic or abiotic stresses, certain elicitors derived from non-self (e.g., flg22, elf18) or altered-self (e.g., SCOOP18) are recognized by their cognate receptors, triggering extracellular alkalinization by inhibiting AHA activity and activating downstream signaling pathways. Mechanistically, using SCOOP18-MIK2 as an example, our findings, along with previous studies, indicate that SCOOP18 is specifically induced in response to CWD^34^ and perceived by its receptor MIK2, leading to the direct phosphorylation of BIK1. BIK1, in turn, phosphorylates GRFs, causing its dissociation from AHAs, resulting in the inhibition of AHA activity and subsequent extracellular alkalinization. This extracellular alkalinization is crucial for signaling activation, likely by promoting ligand-receptor interactions. The BIK1-GRFs-AHAs pathway is activated by multiple elicitors (e.g., flg22, elf18); however, the upstream kinases involved in the phosphorylation relay may differ. For example, in the flg22-FLS2 and elf18-EFR signaling pathways, the co-receptor BAK1 is required to phosphorylate BIK1 and induce extracellular alkalinization. This model provides a mechanistic framework for how plant cell-surface receptors induce extracellular alkalinization upon ligand perception, and how this alkalinization, in turn, regulates immune activation. Given the conservation of these components across plant species, it is plausible that this module could be widely employed by plants to fine-tune pH changes during growth and defense.

The observation that both loss- and gain-of-function mutations in *AHA* result in reduced immune activation in response to elicitors appears to contradict the model in which apoplastic alkalization promotes PTI signaling (Figures 2A-F and 7), especially considering the constitutively alkaline apoplastic pH observed in *aha* mutants^42^. Interestingly, this apparent contradiction is resolved by the effects of transient AHA manipulation using FC and DCCD (Figures 2G-K and S2G), which underscore the importance of dynamic pH changes, rather than a static pH state, in determining signaling strength. This interpretation aligns with the observed reduction in net H⁺ influx in both *AHA-*VIGS and FC-treated cells, where a transient pH gradient may fail to establish effectively (Figures 2N,O and S2L,M). A similar phenomenon was reported in a recent study, showing that dynamic AHA-mediated modulation of apoplastic pH, rather than fixed pH levels, is critical for *Erysiphe cruciferarum* colonization.^82^ Future studies focused on identifying pH-sensing structural elements within receptor kinases and elucidating the molecular mechanisms underlying pH-dependent immune modulation will be essential to advance our understanding of how pH dynamics fine-tune immune signaling.

Unlike in FLS2 signaling in which BAK1 phosphorylates BIK1,^50^ MIK2 directly phosphorylates BIK1 to regulate H⁺ influx (Figure 3). Notably, BAK1 is not genetically required for SCOOP18-triggered H⁺ influx (Figure S6I), suggesting that BIK1 mediates signaling in extracellular alkalinization and other immune responses (e.g., ROS production, calcium burst)^12-15^ through distinct phosphorylation by MIK2 and BAK1, respectively. However, we and others showed that perception of SCOOP peptides induces the association of MIK2 and its co-receptor BAK1,^12-15,46-48^ prompting inquiry into the role of BAK1 in SCOOP-triggered extracellular alkalinization. Considering BAK1 is not involved in initiating SCOOP18-induced H^+^ influx, it will be interesting to investigate its potential roles in the sustained and termination phases of H^+^ influx.

Phosphorylation plays a critical role in regulating AHA activity.^30,31^ A well-characterized regulatory site is the penultimate threonine residue (Thr^948^ in AHA1 and Thr^947^ in AHA2), whose phosphorylation creates a binding site for GRFs, thereby activating the pump.^30,31^ Phosphorylation at Thr^881^ in AHA1/2 also promotes pump activity but independently of GRF binding.^28^ In contrast, phosphorylation at Ser^931^, and likely Ser^899^, has been linked to inhibition of AHA activity.^29,83^ Notably, flg22 treatment in Arabidopsis induces phosphorylation at Ser^899^ while simultaneously reducing phosphorylation at Thr^947^ and Thr^881^, suggesting the involvement of alternative mechanisms to suppress AHA activity during PTI.^84^ In this study, we identify a novel regulatory mechanism in which BIK1 phosphorylates GRF2 at Thr^211^, triggering its dissociation from AHAs and thereby inhibiting AHA activity (Figure 4). Interestingly, the residual inhibition of the AHA1-GRF2^T211A^ interaction by BIK1 suggests that additional factors or pathways may contribute to BIK1-mediated regulation of the AHA-GRF interaction. Given that BIK1 does not directly phosphorylate AHAs, future studies should investigate whether it modulates AHA phosphorylation through indirect mechanisms. Furthermore, elucidating the interplay between distinct phosphorylation events and how they coordinate to fine-tune AHA activity during PTI will be essential. Phosphorylation of GRFs plays a crucial role for their function.^85-87^ In animals, phosphorylation of 14-3-3 proteins reduces their binding to Raf-1 kinase,^86^ and phosphorylation of 14-3-3ζ disrupts its dimeric structure.^87^ In Arabidopsis, cold-activated PM protein COLD-RESPONSIVE PROTEIN KINASE 1 (CRPK1) phosphorylates GRF6 to fine-tune C-repeat (CRT)-binding factor (CBF)-dependent cold signaling.^68^ Notably, Thr^214^ in GRF6, corresponding to Thr^211^ in GRF2, is a major phosphorylation site targeted by CRPK1.^68^ Given the multiple functional roles of BIK1,^17^ it would be interesting to investigate whether BIK1-mediated phosphorylation of GRFs plays a role in other signaling processes. A recent study demonstrated that ten different GRF isoforms were pulled down by PBL19 and PBL20 in Arabidopsis IP-MS experiments,^88^ suggesting that interactions between RLCKs and GRFs are common. Considering the conservation of phosphorylation sites in GRFs, it would be valuable for future studies to systematically investigate whether specific interactions and phosphorylation events between GRFs and RLCKs occur *in planta* in response to various stresses, thereby ensuring proper signaling activation and potentially underlying signaling specificity.

## ACKNOWLEDGMENTS

We thank Jeffrey Leung for providing published plant materials, Libo Shan for VIGS constructs, and Jörg Stelling for yeast strains. We acknowledge all past and current members of the Zipfel group for fruitful discussions. Special thanks to Álvaro D. Fernández-Fernández for confocal imaging and Ryan Toth for critical review of the manuscript prior to submission. This work was supported by the University of Zurich (C.Z.), the Gatsby Charitable Foundation (C.Z. and F.L.H.M.), the European Research Council under the European Union’s Horizon 2020 research and innovation programme no. 773153 (project ‘IMMUNO-PEPTALK’) (C.Z.) and the Swiss National Science Foundation grants no. 31003A_182625, 310030_212382 and 320030_228294 (C.Z.).

## AUTHOR CONTRIBUTIONS

K.Z. and C.Z. conceived and designed the experiments. K.Z., P.D., S.Z., S.C., L.W., and K.W.B. generated materials, performed experiments and/or analyzed the data. T. K. provided AHA phospho-antibodies. B.S. and J.-M.Z. generated the *bik1-2* CRISPR mutant. F.L.H.M., K.W.B. and C.Z. supervised the project. C.Z. obtained funding. K.Z. and C.Z. wrote the manuscript with input from all authors.

## DECLARATION OF INTERESTS

The authors declare no competing interests.

### Lead contact

Further information and requests for resources and reagents should be directed to Cyril Zipfel.

### Data availability

The MS data generated during this study have been deposited to the ProteomeXchange Consortium via the PRIDE^92^ partner repository with the dataset identifier PXD063345.

## METHODS

### *Arabidopsis thaliana* and growth conditions

*Arabidopsis thaliana* ecotype Columbia (Col-0) or Landsberg *erecta* (L*er*) was used as wild-type control. Apart from *ost2-1D* (L*er*), all the mutants investigated in this study are in the Col-0 WT background. The Arabidopsis T-DNA insertion line *aha1* (SAIL_1285_D12) was obtained from the Nottingham Arabidopsis Stock Centre (NASC) and confirmed by PCR and RT-PCR using primers listed in Table S4. All seeds were surface sterilized using chlorine gas for 5-6 h, sown on half strength Murashige and Skoog (½ MS) media supplemented with vitamins, 1 % sucrose, and 0.9 % agar, stratified for 2-3 days in the dark at 4 °C, and moved to a growth chamber with conditions 16-h day/8-h night at respectively 22 °C/18 °C. Five-day-old seedlings were transferred into liquid ½ MS medium and incubated for several days as indicated for the respective experiments. Ten-day-old seedlings were transplanted into soil and grown in a controlled environment growth chamber at 150 µmol light intensity, 60 % relative humidity and 20 °C in a 10-h light cycle.

### Nicotiana benthamiana and growth conditions

*N. benthamiana* plants were grown in a controlled environment chamber at 120 µmol light intensity, 45-60 % relative humidity and 19-21 °C in a 12-h light cycle.

### Microbial strains

*Escherichia coli* DH10β was cultured at 37 °C in LB medium for plasmids extraction. *Agrobacterium tumefaciens* GV3101 was cultured at 28 °C in LB medium for plant transformation. *E. coli* Rosetta (DE3) was cultured at 18 °C in LB medium for protein expression. *Saccharomyces cerevisiae* strain FRY70 was cultured at 28 °C in YPD medium.

### Plasmid construction and generation of transgenic plants

Polymerase chain reaction (PCR) products were amplified from plant DNA, cDNA or plasmid templates using primers listed in Table S4. Point mutations were generated using site-directed mutagenesis primers listed in Table S4. The vector backbones for transient expression in *N. benthamiana* and split-ubiquitin yeast-two-hybrid were generated using Golden Gate system^93^. For transient expression vectors in *N. benthamiana*, AHA1 and AHA2 CDS were cloned into a modified LIIβ F carrying a HA cassette using InFusion. For protein expression vectors in *E. coli*, CDS of MIK2-CD, MIK2*-CD, BAK1-CD, GRF2, GRF2^variants^, GRF10, GRF10^S209A^ and different fragments of AHA1, AHA2 and AHA11 were cloned into pOPINM using InFusion. CDS of BIK1, BIK1* and variants of BIK* were cloned into pET28a using InFusion. CDS of MIK2 was cloned into pGEX-4T-1P using InFusion. For SLC assays *N. benthamiana*, CDS of BAK1, MIK2, MIK2*, EFR, FLS2, AHA1, AHA2 and GRF2 were cloned into a modified pCAMBIA1300 carrying a Nluc part using InFusion. CDS of BIK1, BIK1*, variants of BIK1, AHA1, AHA2 and AHA11 were cloned into a modified pCAMBIA1300 carrying a Cluc part using InFusion. For Agrobacterium-mediated virus-induced gene silencing (VIGS) in Arabidopsis, the specific regions of AHA1, AHA2 and/or AHA11 were cloned into pTRV-RNA2 using InFusion. For yeast-two-hybrid assays, MIK2-Cub, FLS2-Cub, EFR-Cub, AHA2-Nub and AHA11-Nub were cloned into pYeast CEN F2-3 and pYeast CEN F1-2 respectively, using InFusion. AHA1-Nub and MID2-Nub were cloned into pYeast CEN F1-2 using Golden Gate system. For protoplasts assays, CDS of LTI6b, AHA1, GRF2, variants of GRF2 and BIK1^WT/*/Y250F^ were cloned into modified pUC19 vectors with the corresponding tags using Golden Gate system. For generating transgenic plants, the AHA1 promoter and AHA1 CDS were cloned into a modified LIIIβ fin carrying a GFP cassette using InFusion. GRF2 and its variants were cloned into pICSL86944OD using Golden Gate system. Two sgRNAs each for BIK1 and PBL1 were cloned into Level 2 CRISPR vector pICSL4723 using Golden Gate system. The sequences of all genes or promoters were verified by Sanger sequencing. The binary plasmids were introduced into Arabidopsis via *Agrobacterium tumefaciens*-mediated (strain GV3101) transformation by floral dipping^94^.

### RNA analysis

Total RNA was isolated from Arabidopsis seedlings or leaves by FavorPrep Plant Total RNA Purification Mini Kit (Favorgen) following the manufacturer’s instruction. For RT-qPCR analysis, RNA was reverse transcribed to synthesize first-strand cDNA with RevertAid First Strand cDNA Synthesis Kit (Thermo Fisher Scientific) according to the manufacturer’s protocol. RT-qPCR analysis was carried out using PowerUp SYBR Green (Applied Biosystems) with the primers provided in Table S4. Data were analyzed using the 2^-ΔΔCT^ method and normalized to the expression of *ACTIN2*.

### ROS measurement

ROS burst assays were conducted as previously described.^12^ Briefly, leaf discs were harvested from 3- to 4-week-old Arabidopsis plants and floated on 100 μL distilled water overnight or treated as indicated in the figure legends in white 96-well-plates (Greiner Bio-One). Following the specified pretreatment (or in its absence), the water or solution was replaced with ROS assay solution (100 μM luminol, 20 μg mL^-1^ horseradish peroxidase) with the addition of elicitors or ISX, as detailed in the figure legends. For Pep1-triggered ROS measurement, 0.5 μM L-012 was used in place of luminol. Immediately after the addition of the assay solution, luminescence was measured from the plate using a HIGH-RESOLUTION PHOTON COUNTING SYSTEM (HRPCS218, Photek). For kinetic analyses, RLUs were recorded at 1-min intervals over the time periods. *T*_max_ RLU was defined as the interval with the highest recorded value.

### Transient expression

For transient expression in *N. benthamiana*, leaves of four-week-old plants were infiltrated with *A. tumefaciens* strain GV3101 carrying the corresponding constructs as indicated in figure captions. *A. tumefaciens* GV3101 carrying a p19 RNA silencing suppressor construct was co-infiltrated. *N. benthamiana* leaves were collected 2-3 days post-infiltration.

For transient expression in protoplasts, Arabidopsis mesophyll protoplasts were isolated and transfected as previously described,^95^ incubated 8-12 h, and then treated with water or elicitors before collection.

### Split ubiquitin-based membrane yeast two-hybrid (MYTH) assay

The split-ubiquitin yeast two-hybrid assays were performed as previously described^96^ with some modifications. Briefly, the reporter strain FRY70 was constructed by integrating the plasmid FRP795, enabling fluorescent screening through a flow cytometer. MIK2, EFR or FLS2 with C-terminal Cub-LexA-VP16 fusion was used as the bait. AHA1, AHA2 or AHA11 with C-terminal Nub fusion were used as preys. Yeast membrane protein MID2 with C-terminal Nub fusion was used as the negative control. Transformed FRY70 strains carrying the desired constructs were cultured in yeast medium overnight, followed by fluorescence measurement using flow cytometer BD FACSCanto™ II with High Throughput Sampler. FlowJo software was used for data analysis, and the fluorescence intensity median was used to indicate interaction strength.

### Split-luciferase (SLC) assay

*A. tumefaciens* strain GV3101 carrying the desired constructs were infiltrated into leaves of four-week-old *N. benthamiana*. Leaf discs were taken 2-3 days post-infiltration, incubated with 100 µL water containing 1 mM luciferin in a 96-well plate for 15-20 min, and then luminescence was captured with the Spark plate reader. For SCOOP18 treatment, leaves were infiltrated with 1 µM SCOOP18 peptide or water using a needleless syringe for 10-15 min before collection. Equal expression of proteins was verified by western blot.

### Protein extraction and Co-immunoprecipitation (Co-IP) assay

For total protein extraction and phosphorylation activation assay, plant tissues were ground into fine powder in liquid nitrogen, homogenized in protein extraction buffer (50□mM Tris-HCl pH 7.5, 100 mM NaCl, 10 % glycerol, 2 mM EDTA, 5 mM DTT, 1 % IGEPAL CA630, 2 mM Sodium molybdate, 1 mM Sodium fluoride, 1 mM Sodium orthovanadate, 4 mM Sodium tartrate, protease inhibitor cocktail). Samples were incubated at 4 °C for 0.5-1 h and then centrifuged at 16,000 g for 10□min at 4□°C to remove debris. Supernatants were collected with SDS loading buffer for protein gel blot analysis. For Co-IP, supernatants were incubated with GFP-Trap Agarose, RFP-Trap Agarose or anti-FLAG M2 Affinity Gel for 4 h at 4 °C for immunoprecipitation. Following incubation, beads were washed three times with extraction buffer before adding SDS loading buffer. Proteins were separated by SDS-PAGE, transferred to a PVDF membrane, blocked, and probed in TBST-5 % non-fat milk using the corresponding antibodies. Antibodies against the following proteins were used: HA (1:2,500), GFP (1:1,500), RFP (1:1,500), Flag (1:5,000), GST (1:5,000), MBP (1:5,000), His (1:2,000), Luciferase (1:2,000), p44/42 MAPK (1:1,000), BIK1 (1:2,000), BAK1 (1:10,000), BAK1^S612^ (1:2,000), AHA2 (1:2,500), FKBP12 (1:1000), Anti-rabbit IgG-peroxidase (1:10,000) and Anti-mouse IgG-peroxidase (1:15,000).

### Biotin-elf24-EFR interaction assay

Pull-down assays were performed using neutravidin beads under the specified pH conditions. EFR-GFP or FLS2-GFP proteins were extracted from the respective Arabidopsis transgenic plants using extraction buffer (25 mM NaH_2_PO_4_, 25 mM Na_2_HPO_4_, 150 mM NaCl, 1 mM DTT, 0.1 % NP-40, protease cocktail inhibitor). The pH was adjusted by mixing different volume of NaH_2_PO_4_ and Na_2_HPO_4_. Biotin-elf24 was incubated with neutravidin beads in PBS buffer for 1 h at 4 °C, followed by three washes with PBS. The beads were subsequently incubated with EFR-GFP or FLS2-GFP extracts at different pH conditions for 3 h at 4°C. Following incubation, the beads were washed three times with extraction buffer before the addition of SDS loading buffer. Samples were resolved by SDS-PAGE, transferred to PVDF membranes, and detected by immunoblotting with GFP antibody.

### Recombinant protein expression and purification

All recombinant proteins used in this study were expressed in *E. coli* strain BL21 (DE3) Rosetta pLysS. BAK1-CD, MIK2-CD, GRF2, variants of GRF2, GRF10, different fragments of AHA1, AHA2 and AHA11 were expressed as 6×His-MBP fusion proteins in the pOPINM vector. GST-MIK2-CD was expressed using pGEX-4T-1 vector. BIK1, BIK1* (BIK1^K105A/K106A^) and its variants were expressed as 6×His fusion proteins in the pET-28a vector. All recombinant proteins were purified using HisPur cobalt resin (Thermo) or GST-bind resin (Millipore) for 6×His, 6×His-MBP and GST fusions, respectively.

### *In vitro* pull-down assay

Approximately 6 µg each of bait proteins were captured with corresponding resin (GST-bind resin for GST-MIK2-CD or HisPur cobalt resin for His-BIK1), washed three times, then mixed with prey proteins to 2 mL of final volume in buffer (25 mM Tris-HCl pH 7.4, 100 mM NaCl, 0.2 % Triton X-100, 1 mM DTT). Samples were incubated at 4 °C for 1-2 h. The resin was washed four times with buffer and enriched proteins were eluted with 50 µL of SDS-loading buffer. Samples were separated by SDS-PAGE, transferred to PVDF and visualized by blotting with the corresponding antibody.

### *In vitro* kinase assay

Approximately 1 µg of kinase protein was incubated with 1 µg of substrate protein in kinase buffer (25 mM Tris-HCl pH 7.4, 5 mM MnCl_2_, 5 mM MgCl_2_, 1 mM DTT). Reactions were initiated by adding 5 µM ATP along with 0.5 µCi □-^32^P-ATP in a final volume of 30 µL, and incubated at room temperature for 30 min. Reactions were stopped by the addition of SDS-loading buffer and heating at 75 °C for 10 min. Proteins were separated by SDS-PAGE followed by transfer to PVDF membrane and staining with Coomassie brilliant blue G-250. Autoradiographs were imaged using an Amersham Typhoon Phosphorimager (GE Healthcare). For analysis by mass-spectrum, *in vitro* kinase assays were conducted as above, excluding the addition of □-^32^P-ATP.

### Histological staining assay

Root lignification assays were performed as previously described.^34^ Briefly, seven-day-old seedlings grown in the liquid culture were replaced with new liquid media containing either DMSO (mock) or 0.6 μM ISX overnight, and then the seedlings were harvested in 70 % ethanol and stained for lignification using phloroglucinol-HCl. For quantification of lignin deposition in the root, pictures were taken with Lecia microscope. Phloroglucinol-stained areas were quantified with ImageJ.

HPTS staining was performed as previously described.^36^ Briefly, five-day-old seedlings grown in liquid ½ MS media were treated with different elicitors or ISX for the indicated time, followed by supplementation with 1 mM HPTS for 1 h before observation with confocal microscopy (Leica SP5).

### Agrobacterium-mediated virus-induced gene silencing (VIGS)

VIGS assays were performed as previously described.^97^ Briefly, *A. tumefaciens* strain GV3101 carrying the binary TRV vectors *pTRV-RNA1* and *pTRV-RNA2* derivatives, *pTRV-RNA2-GFP* (negative control), *pTRV-RNA2-CLA1* (technical positive control), *pTRV-RNA2-AHA2*, *pTRV-RNA2-AHA2-AHA11*, *pTRV-RNA2-AHA1-AHA2-AHA11* were first cultured in LB medium and then sub-cultured in fresh LB medium containing 10 mM MES and 20 μM acetosyringone overnight. Cells were pelleted and re-suspended in infiltration buffer (10 mM MgCl_2_, 10 mM MES and 200 μM acetosyringone), adjusted to OD600 = 2.0, and incubated at room temperature for 3 h. Bacterial cultures containing pTRV-RNA1 and pTRV-RNA2 derivatives were mixed at a 1:1 ratio and infiltrated into the leaves of two-week-old soil-grown Arabidopsis using a needleless syringe.

### H^+^ flux measurement

Net H^+^ fluxes were measured using non-invasive micro-test technology (NMT) (Younger USA, LLC, MA, USA) with imFluxes V 3.0 software (Younger, USA). To measure net H^+^ fluxes from spongy mesophyll cells, four-week-old Arabidopsis leaves were used. The abaxial epidermis was peeled off from the leaves and fixed into a chamber using a double-sided adhesive tape, followed by incubation in bath solution (0.1 mM CaCl_2_, 0.1 mM KCl, pH 6.0) with or without ISX treatment. The bath solution was then refreshed, and H^+^ fluxes measurement were taken following the addition of flg22, elf18, SCOOP18 or no treatment. Microelectrodes were fabricated by pulling borosilicate glass capillaries (GC150T-10; Harvard Apparatus Ltd), baked for 2 h at 220 **°**C, and silanized using N,N-Dimethyltrimethylsilylamine (Sigma, 41716). Electrodes with a tip diameter of 1-2 µm were filled with Hydrogen ionophore I (Sigma, 95291) and backfilled with 15 mM NaCl, 40 mM KH_2_PO_4_, pH 7.0. H^+^ microelectrodes were calibrated with standard solutions. The microelectrode was positioned about 20 μm from a single cell and moved back and forth between 20-60 μm. Ion fluxes were determined as previously described.^98^

### ATP hydrolytic activity

ATP hydrolysis by the PM H^+^-ATPase was measured in a vanadate-sensitive manner as previously described^44^ with some modifications. Briefly, ten-day-old seedlings or protoplasts were treated with mock, flg22, elf18, SCOOP18 or 0.6 μM ISX for the indicated time (2-5 min) and then collected for preparation of microsomal membranes. The microsomal fraction was then resuspended with ATPase reaction buffer. The ATP hydrolytic activity was determined in a vanadate-sensitive way, and the inorganic phosphate released from ATP was measured.

### LC-MS/MS analysis

For phosphorylation site analysis of His-BIK1* by MIK2 kinase and GRF2 by BIK1, samples were separated on a NuPAGE 4-12 % Bis-Tris gel (Invitrogen) and the corresponding band was excised after staining with InstantBlue (Abcam). For AHA1 IP-MS, two-week-old seedlings of Col-0/*AHA1pro::AHA1-GFP* and Col-0/*GFP-LTI6b* were collected for protein extraction. IP beads were resuspended in 50 µL of elution buffer (Invitrogen) and incubated at 80 °C for 8 min. The samples were run into a NuPAGE 4-12 % Bis-Tris gel and the gel lanes were excised after staining with InstantBlue (Abcam). The excised gel portions were cut into smaller pieces and washed three times with 50 % acetonitrile, 50 mM ammonium bicarbonate (50 % AcN/ABC), 30 min each, followed by dehydration in acetonitrile, 10 min. Gel pieces were then reduced with 10 mM DTT for 30 min at 45 °C followed by alkylation with 55 mM chloroacetamide for 20 min at room temperature, and a further three washes with 50 % AcN/ABC, 30 min each. Gel pieces were dehydrated again with acetonitrile before rehydration with 40□µL of trypsin (Pierce Trypsin Protease) working solution (100□ng of trypsin in 50□mM ammonium bicarbonate, 5 % (v/v) acetonitrile). Where required, gel pieces were covered with 50□mM ammonium bicarbonate to a final volume before incubation at 37□°C overnight. Tryptic peptides were extracted from the gel pieces three times in an equal volume of 50 % acetonitrile, 5 % formic acid (Pierce), 30□min each. Extracted peptides were dried in a speed-vac and resuspended in 2 % acetonitrile/0.2 % trifluoroacetic acid (Sigma). A total of three biological replicates were performed for each sample type.

Samples were analyzed using either an Orbitrap Fusion Tribrid mass spectrometer coupled to a U3000 nano-UPLC, or an Orbitrap Eclipse Tribrid mass spectrometer equipped with a FAIMS Pro Duo interface, with each coupled to a U3000 nano-UPLC system (Thermo Fisher Scientific, Hemel Hempstead, UK).

For analysis by Orbitrap Fusion, approximately 35 % of the dissolved peptides were injected onto a reverse phase trap column NanoEase m/z Symmetry C18, beads diameter 5□μm, inner diameter 180□μm□×□20□mm length (Waters). Trap column flow rate was 20□μl□min^-1^ in 2 % acetonitrile, 0.05 % TFA. Peptides were eluted from trap column onto the analytical column NanoEase m/z HSS C18 T3 Column, beads diameter 1.8□μm, inner diameter 75□μm□×□250□mm length (Waters). The column was equilibrated with 3 % B (B, 80 % acetonitrile in 0.05 % formic acid (FA); A, 0.1 % FA) before subsequent elution with the following steps of a linear gradient: 2.5□min 3 % B, 5□min 6.3 % B, 13□min 12.5 % B, 50□min 42.5 % B, 58□min 50 % B, 61□min 65 % B, 63□min 99 % B, 66□min 99 % B, 67□min 3 % B, 90□min 3 % B. The flow rate was set to 200□nl□min^-1^. The mass spectrometer was operated in positive ion mode with nano-electrospray ion source. Molecular ions were generated by applying voltage +2.2□kV to a conductive union coupling the column outlet with fused silica PicoTip emitter, ID 10□μm (New Objective) and the ion transfer capillary temperature was set to 275 °C. The mass spectrometer was operated in data-dependent mode using a full scan, m/z range 300-1800, nominal resolution of 120000, target value 1□×□10^6^, followed by MS/MS scans of the 40 most abundant ions. MS/MS spectra were acquired using normalized collision energy of 30 %, isolation width of 1.6□m/z, resolution of 120000 and a target value set to 1□×□10^5^. Precursor ions with charge states 2-7 were selected for fragmentation and put on a dynamic exclusion list for 30□s. To improve detection of phosphorylation, multistage activation was applied for detection of -98, -49 or -32.7 from the precursor (corresponding to the neutral loss of phosphoric acid from +1, +2 and +3 charge states, respectively) during any of the MS/MS scans. The minimum automatic gain control target was set to 5□×□10^3^ and intensity threshold was calculated to be 4.8□×□10^4^. The peptide match feature was set to the preferred mode and the feature to exclude isotopes was enabled.

For Orbitrap Eclipse Tribrid equipped with FAIMS, approximately 20 % of the digested peptides were loaded onto a trap cartridge (PepMap™ Neo Trap Cartridge, C18, 5 μm, 0.3 ×□5 mm, Thermo) with 0.1 % TFA at 15 μl min^-1^ for 3 min. The trap column was then switched in-line with the analytical column (Aurora Frontier TS, 60 cm nanoflow UHPLC column, ID 75 μm, reversed phase C18, 1.7 μm, 120 Å; IonOpticks, Fitzroy, Australia) for separation at 60 °C using the following gradient of solvents A (water, 0.1 % formic acid) and B (80 % acetonitrile, 0.1 % formic acid) at a flow rate of 0.26 μl min^-1^: 0-3 min 1 % B (parallel to trapping); 3-10 min increase B (curve 4) to 8 %; 10-102 min linear increase B to 48; followed by a ramp to 99 % B and re-equilibration to 0 % B, for a total of 140 min runtime. Mass spectrometry data were acquired with the FAIMS device set to three compensation voltages (-35V, -50V, -65V) at standard resolution for 1.0 s each with the following MS settings in positive ion mode: OT resolution 120 K, profile mode, mass range m/z 300-1600, normalized AGC target 100 %, max inject time 50 ms; MS2 in IT Turbo mode: quadrupole isolation window 1 Da, charge states 2-5, threshold 1e4, HCD CE = 30, AGC target standard, max. injection time dynamic, dynamic exclusion 1 count for 15 s with mass tolerance of ±10 ppm, one charge state per precursor only.

Peak lists in the form of Mascot generic files were prepared from raw data files using MS Convert v.3.0 (Proteowizard) and sent to a peptide search on Mascot server v.2.8.2 using Mascot Daemon (Matrix Science) against an in-house constructs and contaminants database and the *E. coli* K12 protein database. Tryptic peptides with up to two possible mis-cleavages and charge states +2 and +3 were allowed in the search. The following peptide modifications were included in the search: carbamidomethylated cysteine (fixed), oxidized methionine (variable), with phosphorylated serine, threonine and tyrosine (variable). Data were searched with a monoisotopic precursor and fragment ion mass tolerance 10□ppm and 0.8□Da respectively. Decoy database was used to validate peptide sequence matches. Mascot results were combined in Scaffold v.5.0.0 (Proteome Software) and filtered to show only phospho-peptides. For His-BIK1* phosphopeptide analysis, peptide probability and protein threshold were set at 50.0 % and 99.0 % respectively and spectra were manually curated to verify validity of identification and position of the phosphorylated residue. For His-MBP-GRF2 analysis, peptide probability and protein threshold were each set at 1 % false discovery rate (FDR). Spectral counts for each identified phospho-peptide from three biological replicates were summed. Spectral counts from mis-cleaved peptides identifying the same phosphorylation site were then summed to give final spectral counts for each site. For identification of GRFs potentially interacting with AHA1, peptide probability and protein threshold were each set at 1 % FDR. Mean spectral counts from three biological replicates each for AHA1-GFP and GFP-LTI6b controls were then compared.

### Quantification and statistical analysis

Data for quantification analyses are presented as mean ± standard deviation (SD) or standard error (SE). The normal distribution of the data is assessed using the Kolmogorov-Smirnov test. If the data follows a normal distribution, a two-tailed Student’s *t*-test is employed for two-group comparisons, while one-way ANOVA followed by Tukey’s test is used for multiple comparisons. If the data does not follow normal distribution, a two-tailed Mann-Whitney test is applied for two-group comparisons, and a Kruskal-Wallis test followed by Dunn’s test is used for multiple comparisons. Two-way ANOVA is used for statistical analysis of H^+^ influx. *P*-values indicate the data analyzed by two-tailed Student’s *t*-test, two-tailed Mann-Whitney test or two-way ANOVA. Different letters denote statistically significant differences according to one-way ANOVA followed by Tukey’s test or Kruskal-Wallis test followed by Dunn’s test (*P* < 0.05). The number of replicates is shown in the figure legends.

## FIGURE LEGENDS

**Figure S1.**
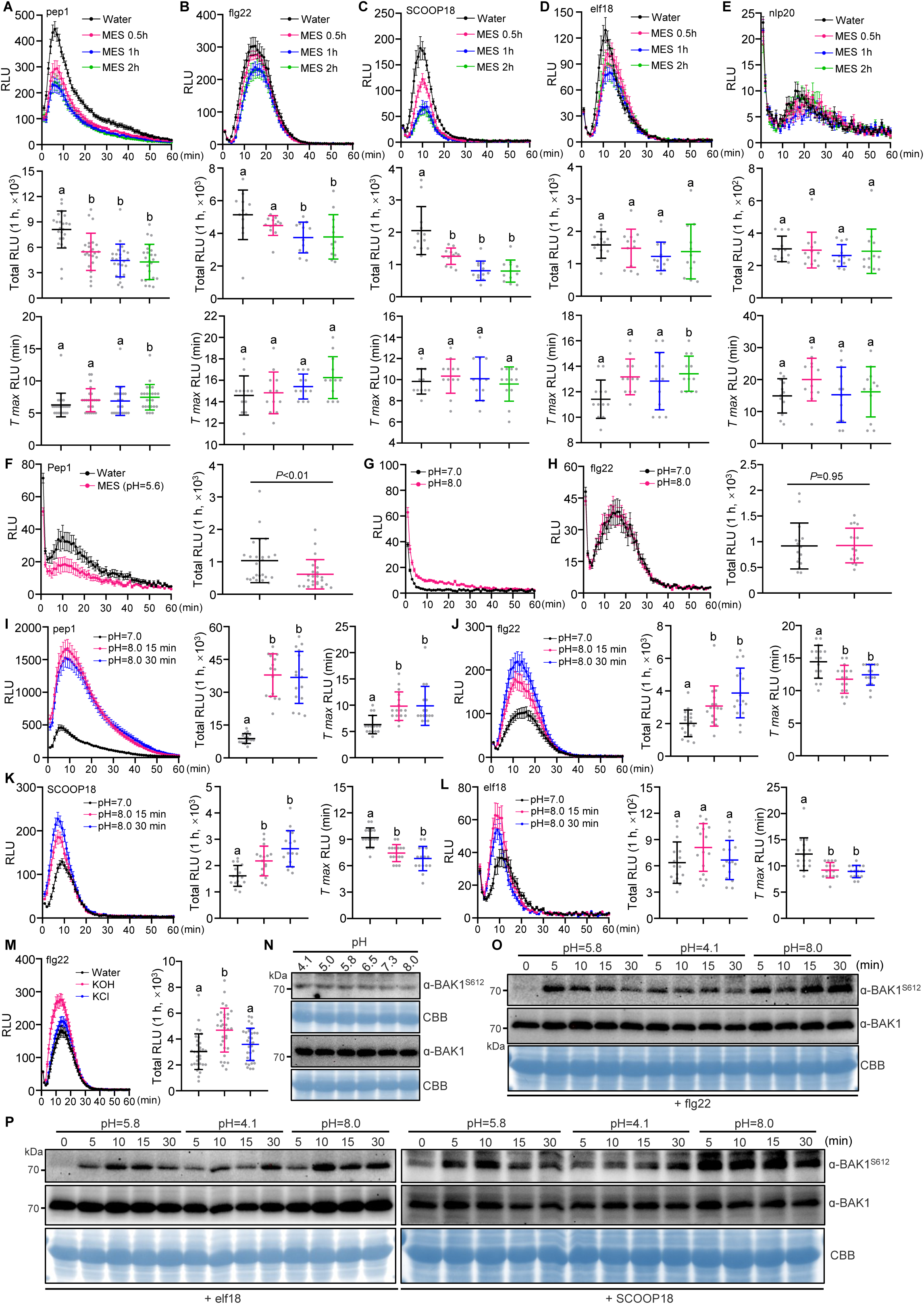
Extracellular alkalinization promotes PTI signaling. (A-E) ROS production after treatment with different elicitors. Leaf discs of Col-0 were pretreated with or without 2 mM MES (pH 5.6) for the indicated time points, followed by treatment with pep1 (A), flg22 (B), SCOOP18 (C), elf18 (D), or nlp20 (E). Values are means of RLU of independent leaf discs (upper panel), total RLU (middle panel) or the time to maximum RLU (*T_max_* RLU) (down panel) over 1 h. The data are shown as mean ± SE (upper panel) or mean ± SD (middle and down panels), n = 24 (A), n = 12 (B-E). (F) ROS production after treatment with Pep1. Leaf discs of Col-0 were pretreated with or without 2 mM MES (pH 5.6) for 24 h, followed by treatment with Pep1. Values are means of RLU of independent leaf discs (left panel) or total RLU (right panel) over 1 h. The data are shown as mean ± SE (left panel) or mean ± SD (right panel), n = 24. (G) ROS production in different pH solutions. Leaf discs were treated with solutions with pH 7.0 or 8.0. Values are means of RLU of independent leaf discs over 1 h ± SE, n = 24. (H) ROS production after treatment with flg22. Leaf discs of Col-0 were pretreated with different pH solutions for 1 min, followed by treatment with flg22. Values are means of RLU of independent leaf discs (left panel) or total RLU (right panel) over 1 h. The data are shown as mean ± SE (left panel) or mean ± SD (right panel), n = 16. (I-L) ROS production after treatment with different elicitors. Leaf discs of Col-0 were pretreated with solutions (pH 7.0 or 8.0) for the indicated time points, followed by treatment with pep1 (I), flg22 (J), SCOOP18 (K) or elf18 (L). Values are means of RLU of independent leaf discs (left panel), total RLU (middle panel) or the time to maximum RLU (*T_max_* RLU) (right panel) over 1 h. The data are shown as mean ± SE (left panel) or mean ± SD (middle and right panels), n = 16. (M) ROS production after treatment with flg22. Leaf discs of Col-0 were pretreated with 0.3 mM KOH (pH 8.0) or 0.3 mM KCl for 30 min, followed by treatment with flg22. Values are means of RLU of independent leaf discs (left panel) or total RLU (right panel) over 1 h. The data are shown as mean ± SE (left panel) or mean ± SD (right panel), n = 32. (N-P) BAK1 phosphorylation in two-week-old seedlings of Col-0. Seedlings pretreated with ½ MS with the indicated pH for 12 h was collected (N) or followed by treatment with flg22 (O), elf18 (P, left panel) or SCOOP18 (P, right panel). 100 nM flg22, 100 nM elf18, 1 μM SCOOP18, 1 μM Pep1 or 1 μM nlp20 was used for the treatment. Different letters in (middle panels of B, C, I and K; D; down panel of E; right panel of J, L and M) denote statistically significant differences according to one-way ANOVA followed by the Tukey’s test (*P* < 0.05). Different letters in (A; down panels of B and C; middle panels of E and J; right panels of I and K) denote statistically significant differences according to Kruskal-Wallis test followed by Dunn’s test (*P* < 0.05). *P*-values indicate the data were analyzed by two-tailed Mann-Whitney test (F) or two-tailed Student’s *t*-test (H). CBB was used as a loading control (N-P). The experiments were repeated at least twice (N-P) or three times (A-M) with similar results.

**Figure S2.**
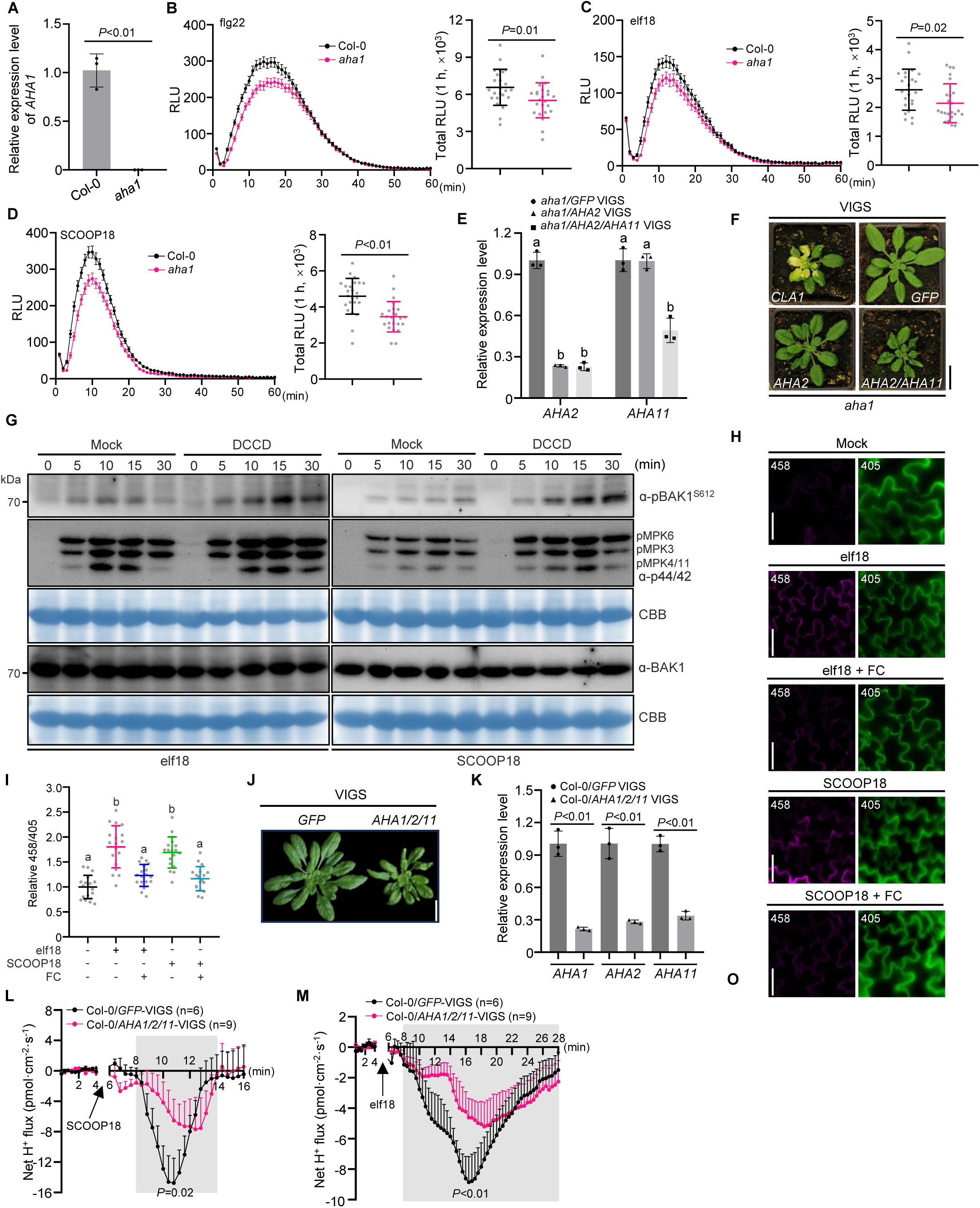
AHAs are essential for elicitor-triggered H^+^ influx, extracellular alkalinization, and immune signaling. (A) RT-qPCR analysis of *AHA1* expression levels in Col-0 and *aha1* T-DNA line. (B-D) ROS production after treatment with different elicitors. Leaf discs of Col-0 and *aha1* were treated with flg22 (B), elf18 (C) or SCOOP18 (D). Values are means of RLU of independent leaf discs (left panel) or total RLU (right panel) over 1 h. The data are shown as mean ± SE (upper panel) or mean ± SD (down panel), n = 24. (E) RT-qPCR analysis of *AHA2* and *AHA11* expression levels in *GFP*-VIGS, *AHA2*-VIGS and *AHA2/AHA11*-VIGS in *aha1* background. (F) Plant phenotypes are shown three weeks after VIGS of *AHA2*, *AHA2/AHA11*, *GFP* (negative control) or *CLOROPLASTOS ALTERADOS 1* (*CLA1*) (technical positive control). Scale bars, 2 cm. (G) BAK1 and MPK kinase phosphorylation in two-week-old seedlings of Col-0. Seedlings pretreated with 10 μM DCCD or mock for 2 h, followed by treatment with elf18 (left panel), or SCOOP18 (right panel). CBB was used as a loading control. (H and I) Confocal images showing the apoplastic pH in cotyledon cells treated with elicitors. Five-day-old seedlings treated with elf18 or SCOOP18, with or without 5 μM FC, for 1 h and stained with HPTS. Changes in pH (imaging, H; quantification, I; mean ± SD, n = 18) were visualized with ratiometric values of fluorescent HPTS. (J) Plant phenotypes are shown three weeks after VIGS of *AHA1/AHA2/AHA11* or *GFP* (negative control) in Col-0. Scale bars, 2 cm. (K) RT-qPCR analysis of *AHA1*, *AHA2* and *AHA11* expression levels in *GFP*-VIGS, *AHA1/AHA2/AHA11*-VIGS in Col-0. (L and M) Net H^+^ flux kinetics upon SCOOP18 and elf18 treatment determined with NMT. Four-week-old mesophyll cells treated with SCOOP18 (L) or elf18 (M). The data are shown as mean ± SE. 100 nM flg22, 100 nM elf18 or 1 μM SCOOP18 was used for the treatment. Different letters in (E and I) denote statistically significant differences according to one-way ANOVA followed by the Tukey’s test (*P* < 0.05). *P*-values indicate the data were analyzed by two-tailed Student’s *t*-test (A, B, D and K), two-tailed Mann-Whitney test (C) or two-way ANOVA (shaded areas of L and M). The experiments were repeated at least twice (A, E, G and K-M) or three times (B-D, H and I) with similar results.

**Figure S3.**
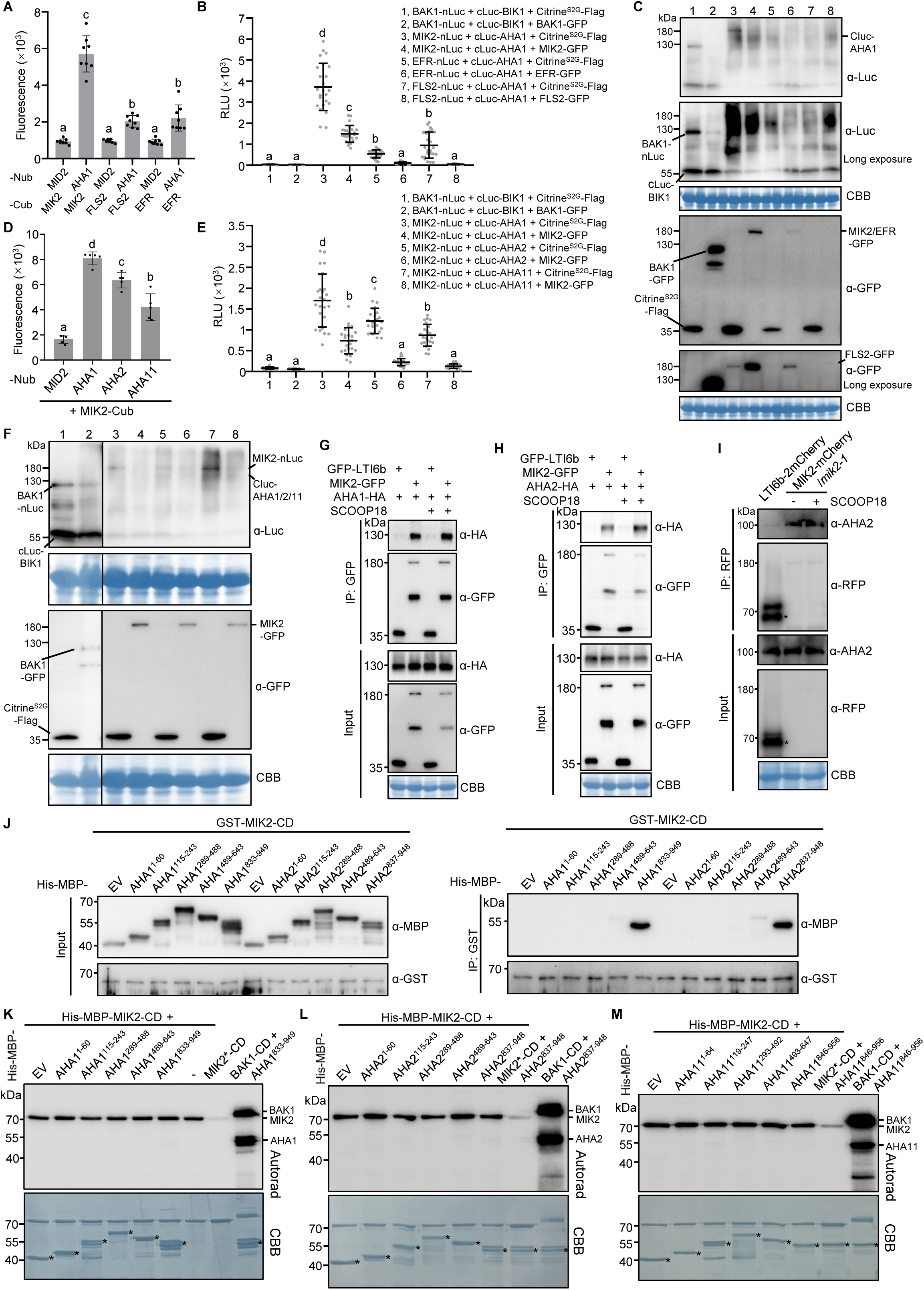
Interaction and phosphorylation analysis of AHAs with immune receptors. (A) Interaction in yeast co-expressing MIK2, FLS2 or EFR-Cub and AHA1-Nub. The yeast plasma membrane protein Mating-Induced Death 2 (MID2) was used as a negative control. (B and C) AHA1 interacts with MIK2, EFR and FLS2 *in N. benthamiana*. SLC assays were conducted with the indicated constructs. BAK1-, MIK2-, EFR- or FLS2-GFP were used for respective interaction competition to show interaction specificity. BAK1-BIK1 was used as a negative control. Values are means of RLU ± SD, n = 24 independent leaf discs (B). Protein expression was detected by western blot (C). (D) Interaction in yeast co-expressing MIK2-Cub and AHA1, AHA2 or AHA11-Nub. The yeast plasma membrane protein MID2 was used as a negative control. (E and F) MIK2 interacts with AHA1, AHA2 and AHA11 in *N. benthamiana*. SLC assays were conducted with the indicated constructs. MIK2-GFP was used for interaction competition to show interaction specificity. BAK1-BIK1 was used as a negative control. Values are means of RLU ± SD, n = 24 independent leaf discs (E). Protein expression was detected by western blot (F). (G and H) Co-immunoprecipitation of MIK2-GFP with AHA1-HA (G) or AHA2-HA (H), transiently expressed in *N*. *benthamiana* leaves and treated with SCOOP18 or mock for 10 min. The plasma membrane marker GFP-LTI6b was used as a negative control. (I) Co-immunoprecipitation of MIK2 and AHA2 in stable transgenic plants *mik2-1/MIK2pro::MIK2-mCherry*. Seedlings were treated with SCOOP18 or mock for 10 min. Endogenous AHA2 protein was detected in immunoblots using α-AHA2 antibody. The LTI6b-2×mCherry transgenic seedlings were used as a negative control. (J) GST pulldown of different fragments of AHA1 and AHA2 fused to His-MBP by GST-MIK2-CD. The empty vector (EV), His-MBP, was used as a negative control. (K-M) Autoradiogram of *in vitro* kinase assay using MIK2-CD with different fragments of AHA1 (K), AHA2 (L) or AHA11 (M). BAK1-CD and His-MBP were used as positive and negative controls, respectively. CBB indicates loading control (C, F-I) or loading of the protein (K-M). Different letters in (A, B, D and E) denote statistically significant differences according to one-way ANOVA followed by the Tukey’s test (*P* < 0.05). The experiments were repeated at least twice (C, F and J-M) or 3 times (A, B, D, E, G-I) with similar results.

**Figure S4.**
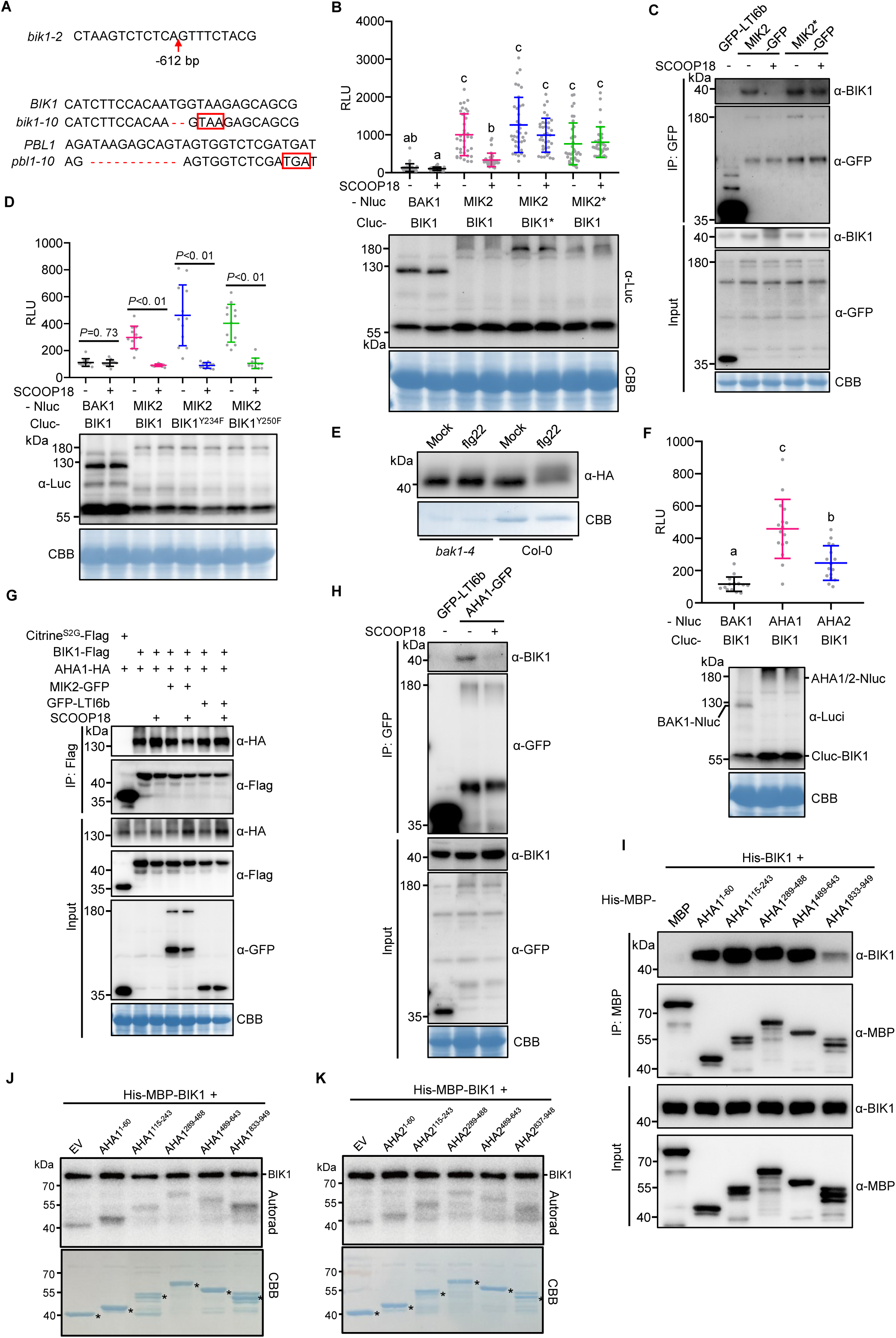
Analysis of MIK2-BIK1 and BIK1-AHA1/2 interactions and phosphorylation. (A) Mutation forms in *bik1-2* and *bik1-10 pbl1-10* mutants. Red arrows and dash lines indicate deletions and red boxes indicate early stop codon. (B and D) MIK2-BIK1 interaction in *N. benthamiana*. SLC assays were conducted with the indicated constructs in the presence or absence of 1 μM SCOOP18 for 10 min. Protein expression was detected by western blot (down panel). BAK1-BIK1 was used as a negative control. Values are means of RLU of independent leaf discs ± SD, n = 36 (B), n = 12 (D). (C) Co-immunoprecipitation of MIK2 and BIK1 in stable transgenic plants expressing MIK2-GFP or MIK2*-GFP (MIK2^D905N^-GFP). Seedlings were treated with 1 μM SCOOP18 or mock for 10 min. Endogenous BIK1 protein was detected in immunoblot using α-BIK1 antibody. The GFP-LTI6b transgenic seedlings was used as a negative control. (E) flg22-induced BIK1 mobility shift in protoplasts. Protoplasts from Col-0 or *bak1-4* were transfected with BIK1-2×HA and then treated with 100 nM flg22 for 10 min. (F) BIK1 interacts with AHA1 and AHA2 in *N. benthamiana*. SLC assays were conducted with the indicated constructs. Protein expression was detected by western blot (down panel). BAK1 was used as negative control. Values are means of RLU ± SD, n = 16. (G) Co-immunoprecipitation of AHA1-HA and BIK1-Flag transiently expressed in *N*. *benthamiana* leaves treated with 1 μM SCOOP18 or mock for 10 min. The plasma membrane Citrine^S2G^-Flag and GFP-LTI6b were used as negative controls. (H) Co-immunoprecipitation of AHA1 and BIK1 in stable transgenic plants Col-0/*AHA1pro::AHA1-GFP*. Seedlings were treated with 1 μM SCOOP18 or mock for 10 min. Endogenous BIK1 protein was detected in immunoblots using α-BIK1 antibody. The GFP-LTI6b transgenic seedlings were used as a negative control. (I) MBP pulldown of His-BIK1 by different fragments of AHA1 fused to His-MBP. The His-MBP-MBP was used as a negative control. (J and K) Autoradiogram of *in vitro* kinase assay using BIK1 with different fragments of AHA1 (J) or AHA2 (K). These experiments were performed together with Figure 3H, which served as a positive control. The asterisks denote the corresponding proteins. CBB indicates loading control (B-H) or loading of the protein (J and K). Different letters in (B) and (F) indicate statistically significant differences (*P* < 0.05), determined by one-way ANOVA with Tukey’s test (F) or Kruskal-Wallis test with Dunn’s test (B). *P*-values indicate the data were analyzed by two-tailed Student’s *t*-test (D). The experiments were repeated at least twice (E, G and I-K) or three times (B-D, F and H) with similar results.

**Figure S5.**
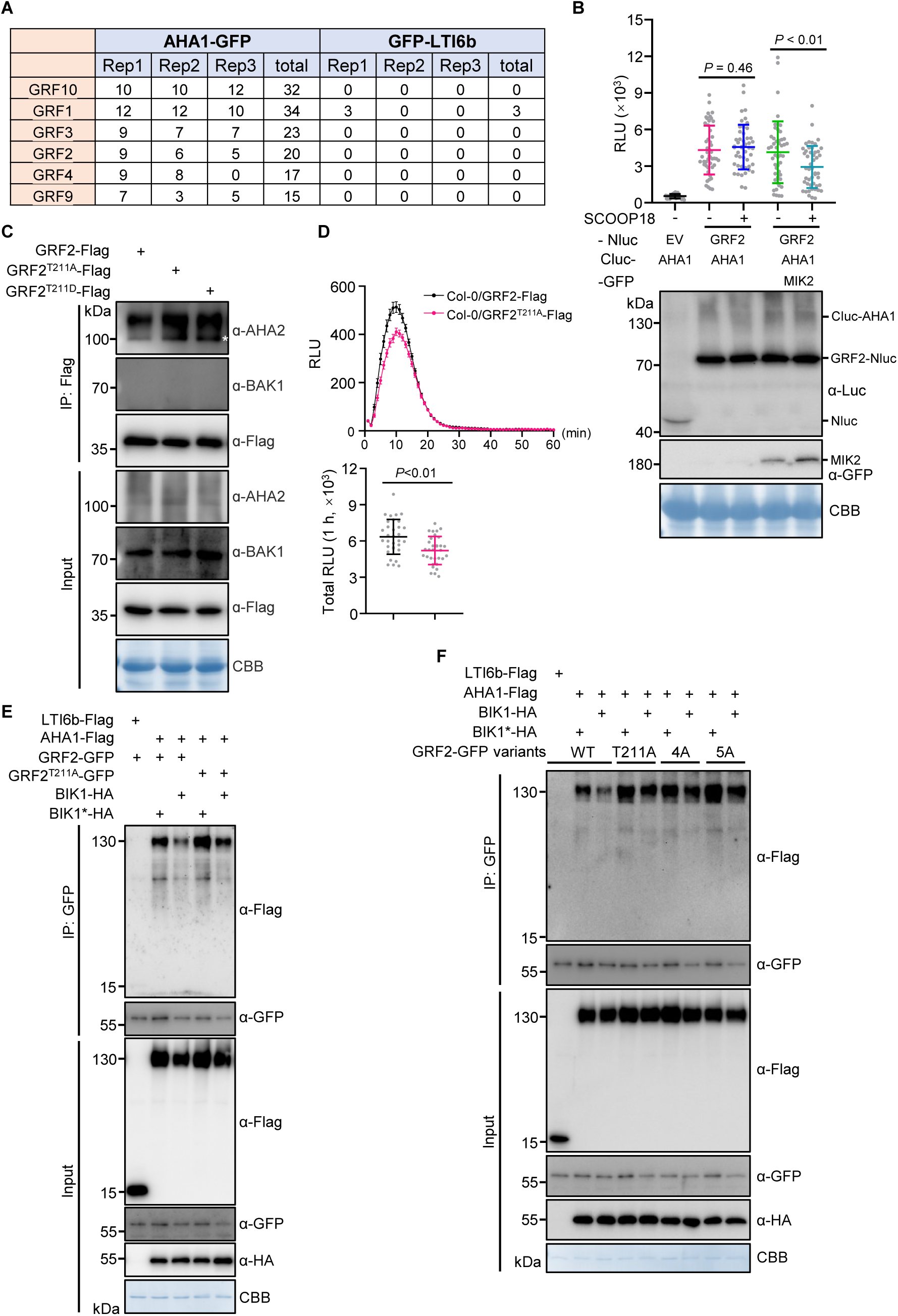
Phosphorylation of GRF2 at Thr211 is required for its dissociation from AHA1 and for immune signaling. (A) The total spectrum counts of GRFs identified in three independent immunoprecipitation assays detected by HPLC-MS/MS. Two-week-old seedlings of Col-0/*AHA1pro::AHA1*-GFP or Col-0/*35S::GFP-LTI6b* were collected for protein extraction and immunoprecipitation. (B) AHA1-GRF2 interaction after SCOOP18 treatment in *N*. *benthamiana*. SLC assays were conducted with the indicated constructs. Protein expression was detected by western blot (down panel). The empty vector (Nluc) was used as a negative control. Values are means of RLU of independent leaf discs ± SD, n = 32 or 48. (C) Co-immunoprecipitation of GRF2 variants (GRF2-Flag, GRF2^T211A^-Flag, and GRF2^T211D^-Flag) with AHA2 in stable transgenic plants. AHA2 protein was detected by immunoblotting using the α-AHA2 antibody, while endogenous BAK1 served as a negative control. The asterisk denotes the endogenous AHA2. (D) ROS production after treatment with SCOOP18. Leaf discs of Col-0/*35S::GRF2-Flag* and Col-0/*35S::GRF2^T211A^-Flag* were treated with 1 μM SCOOP18. Equal protein expression was confirmed, as shown in the input of Figure S5C. Values are means of RLU of independent leaf discs (upper panel) or total RLU (down panel) over 1 h. The data are shown as mean ± SE (upper panel) or mean ± SD (down panel), n = 32. (E and F) Effect of BIK1 on the interaction between AHA1 and different GRF2 variants using Co-IP assays. Protoplasts from Col-0 were transfected with the indicated constructs. The LTI6b-Flag was used as a negative control. CBB indicates loading control (B, C, E and F). *P*-values indicate the data were analyzed by two-tailed Mann-Whitney test (B) or two-tailed Student’s *t*-test (D). The experiments were repeated at least twice (C) or three times (A, B and D-F) with similar results.

**Figure S6.**
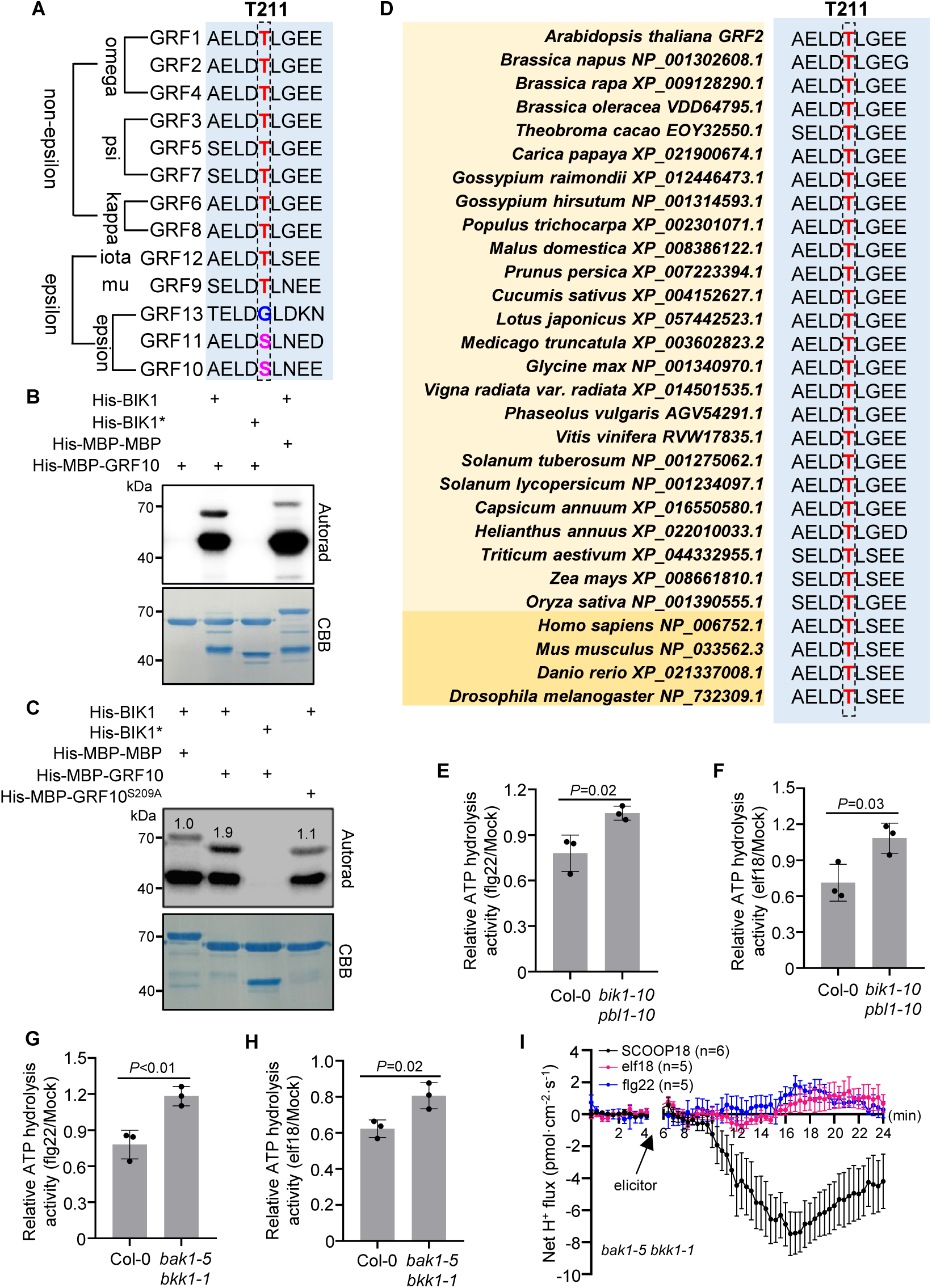
Conservation analysis of BAK1, BIK1 and Thr^211^ of GRF2 in AHA activity regulation. (A) Protein sequence alignment of Arabidopsis GRF family members. Partial amino acid sequences of GRF1 to GRF13 were shown, with the Thr^211^ residue in GRF2 and the corresponding residues in other GRF proteins highlighted and boxed. (B) BIK1 phosphorylates GRF10 *in vitro*. Autoradiogram of *in vitro* kinase assay was conducted with the indicated protein combination. His-MBP-MBP was used as a negative control. (C) *In vitro* phosphorylation assays for analyzing Ser^209^ in GRF10. The assays were conducted with the indicated protein combination. His-MBP-MBP was used as a negative control. Relative ratio of ^32^P incorporation (Autorad/CBB) was quantified using ImageJ and is indicated above the bands. (D) GRF2^T211^ is a conserved site across kingdoms. Sequence analysis of the residues surrounding GRF2 Thr^211^ in 14-3-3 proteins from various cross-kingdom species is shown. The conserved residue is highlighted in red and boxed. (E and F) Quantification of ATP hydrolysis activity in Col-0 and *bik1-10 pbl1-10*. Ten-day-old seedlings treated with flg22 (E) or elf18 (F) for 5 min and then collected for ATP hydrolysis activity measurement. (G and H) Quantification of ATP hydrolysis activity in Col-0 and *bak1-5 bkk1-1*. Ten-day-old seedlings treated with flg22 (G) or elf18 (H) for 5 min and then collected for ATP hydrolysis activity measurement. Experiments shown in Figure S6E and Figure S6G were performed together and share the Col-0 data. (I) Net H^+^ flux kinetics upon flg22, elf18 and SCOOP18 treatment determined with NMT. Four-week-old *bak1-5 bkk1-1* mesophyll cells treated with 100nM flg22, 100nM elf18 or 1 μM SCOOP18. The data are shown as mean ± SE. CBB indicates loading of the protein (B and C). *P*-values indicate the data were analyzed by two-tailed Student’s *t*-test (E-H). The experiments were repeated at least twice (B and C) or three times (E-I) with similar results.

**Figure S7.**
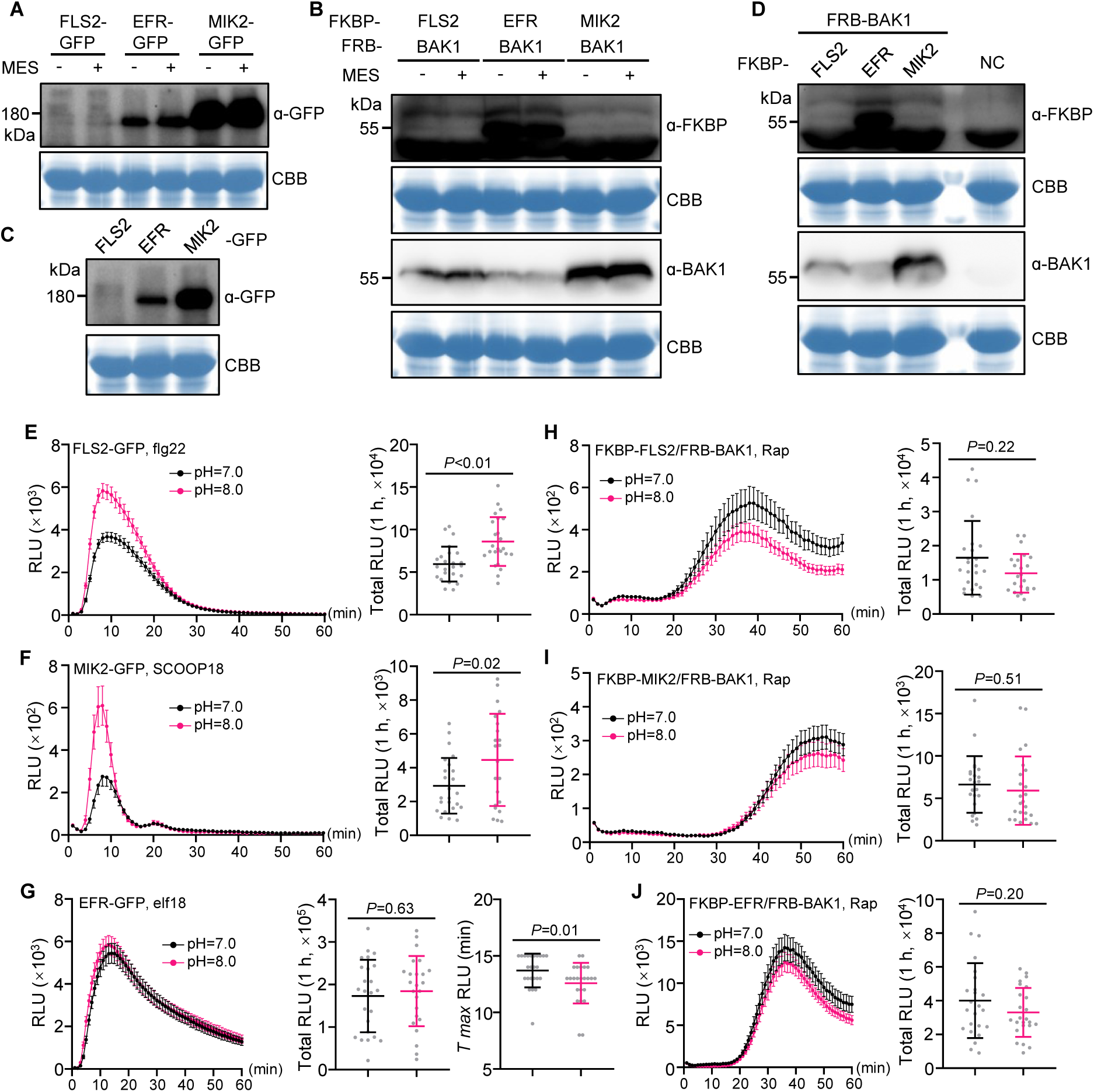
Extracellular alkalinization promotes PTI signaling via the extracellular domains of receptors. (A-D) Protein expression in *N. benthamiana* for Figures 5A-5L was detected by western blot. Figures 5A-5C (A), Figures 5D-5F (B), Figures 5G-5I (C) and Figures 5J-5L (D). Non-infiltrated leaf discs served as the negative control (NC) for D. CBB was used as a loading control. (E-G) ROS production in *N. benthamiana* after treatment with different elicitors. Leaf discs of *N*. *benthamiana* leaves transiently expressing FLS2, MIK2, or EFR were pretreated with solutions with pH 7.0 or 8.0 for 30 min, followed by treatment with 100 nM flg22 (E), 1 μM SCOOP18 (F) or 100 nM elf18 (G). Values are means of RLU of independent leaf discs (left panel), total RLU (right panels of E and F; middle panel of G) or *T_max_* RLU (right panel of G) over 1 h. The data are shown as mean ± SE (left panel) or mean ± SD (total RLU and *T_max_* RLU), with n = 24, except for data from pH = 8.0 in F, where n = 22. (H-J) ROS production in *N. benthamiana* induced by Rap. Leaf discs of *N*. *benthamiana* leaves transiently expressing intracellular domains of FLS2/BAK1, MIK2/BAK1, or EFR/BAK1 were pretreated with solutions with pH 7.0 or 8.0 for 30 min, followed by treatment with 1 μM Rap. Values are means of RLU of independent leaf discs (left panel) or total RLU (right panel) over 1 h. The data are shown as mean ± SE (left panel) or mean ± SD (right panel), with n = 24, except for data from pH = 8.0 in H, where n = 23. *P*-values indicate the data were analyzed by two-tailed Student’s *t*-test (E-G, I and J) or two-tailed Mann-Whitney test (H). The experiments were repeated at least twice (A-D) or three times (E-J) with similar results.

## REFERENCES

1. Couto, D., and Zipfel, C. (2016). Regulation of pattern recognition receptor signalling in plants. Nat. Rev. Immunol. 16, 537–552.

2. Dievart, A., Gottin, C., Perin, C., Ranwez, V., and Chantret, N. (2020). Origin and diversity of plant receptor-like kinases. Annu. Rev. Plant Biol. 71, 131–156.

3. Tang, D., Wang, G., and Zhou, J.M. (2017). Receptor kinases in plant-pathogen interactions: more than pattern recognition. Plant cell 29, 618–637.

4. Escocard de Azevedo Manhaes, A.M., Ortiz-Morea, F.A., He, P., and Shan, L. (2021). Plant plasma membrane-resident receptors: surveillance for infections and coordination for growth and development. J. Integr. Plant Biol. 63, 79–101.

5. Hohmann, U., Lau, K., and Hothorn, M. (2017). The structural basis of ligand perception and signal activation by receptor kinases. Annu. Rev. Plant Biol. 68, 109–137.

6. Gómez-Gómez, L., and Boller, T. (2000). FLS2: an LRR receptor-like kinase involved in the perception of the bacterial elicitor flagellin in Arabidopsis. Mol. Cell 5, 1003–1011.

7. Zipfel, C., Kunze, G., Chinchilla, D., Caniard, A., Jones, J.D., Boller, T., and Felix, G. (2006). Perception of the bacterial PAMP EF-Tu by the receptor EFR restricts Agrobacterium-mediated transformation. Cell 125, 749–760.

8. Gust, A.A., Pruitt, R., and Nurnberger, T. (2017). Sensing danger: key to activating plant immunity. Trends Plant Sci. 22, 779–791.

9. Tanaka, K., and Heil, M. (2021). Damage-associated molecular patterns (DAMPs) in plant innate immunity: applying the danger model and evolutionary perspectives. Annu. Rev. Phytopathol. 59, 53–75.

10. Huffaker, A., Pearce, G., and Ryan, C.A. (2006). An endogenous peptide signal in Arabidopsis activates components of the innate immune response. Proc. Natl. Acad. Sci. USA 103, 10098–10103.

11. Yamaguchi, Y., Pearce, G., and Ryan, C.A. (2006). The cell surface leucine-rich repeat receptor for AtPep1, an endogenous peptide elicitor in Arabidopsis, is functional in transgenic tobacco cells. Proc. Natl. Acad. Sci. USA 103, 10104–10109.

12. Rhodes, J., Yang, H., Moussu, S., Boutrot, F., Santiago, J., and Zipfel, C. (2021). Perception of a divergent family of phytocytokines by the Arabidopsis receptor kinase MIK2. Nat. Commun. 12, 705.

13. Hou, S., Liu, D., Huang, S., Luo, D., Liu, Z., Xiang, Q., Wang, P., Mu, R., Han, Z., Chen, S., et al. (2021). The Arabidopsis MIK2 receptor elicits immunity by sensing a conserved signature from phytocytokines and microbes. Nat. Commun. 12, 5494.

14. Yang, H., Kim, X., Sklenar, J., Aubourg, S., Sancho-Andres, G., Stahl, E., Guillou, M.C., Gigli-Bisceglia, N., Tran Van Canh, L., Bender, K.W., et al. (2023). Subtilase-mediated biogenesis of the expanded family of SERINE RICH ENDOGENOUS PEPTIDES. Nat. Plants 9, 2085–2094.

15. Zhang, J., Zhao, J., Yang, Y., Bao, Q., Li, Y., Wang, H., and Hou, S. (2022). EWR1 as a SCOOP peptide activates MIK2-dependent immunity in Arabidopsis. J. Plant Interact. 17, 562–568.

16. Bender, K.W., and Zipfel, C. (2023). Paradigms of receptor kinase signaling in plants. Biochem. J. 480, 835–854.

17. DeFalco, T.A., and Zipfel, C. (2021). Molecular mechanisms of early plant pattern-triggered immune signaling. Mol. Cell 81, 3449–3467.

18. Liang, X., and Zhou, J.M. (2018). Receptor-like cytoplasmic kinases: central players in plant receptor kinase-mediated signaling. Annu. Rev. Plant Biol. 69, 267–299.

19. Yu, X., Feng, B., He, P., and Shan, L. (2017). From chaos to harmony: responses and signaling upon microbial pattern recognition. Annu. Rev. Phytopathol. 55, 109–137.

20. Felix, G., Regenass, M., and Boller, T. (1993). Specific perception of subnanomolar concentrations of chitin fragments by tomato cells: induction of extracellular alkalinization, changes in protein phosphorylation, and establishment of a refractory state. Plant J. 4, 307–316.

21. Felix, G., Duran, J.D., Volko, S., and Boller, T. (1999). Plants have a sensitive perception system for the most conserved domain of bacterial flagellin. Plant J. 18, 265–276.

22. Felix G, G.D., Regenass M, Boller T. (1991). Rapid changes of protein phosphorylation are involved in transduction of the elicitor signal in plant cells. Proc. Natl. Acad. Sci. USA. 88, 8831–8834.

23. Kunze, G., Zipfel, C., Robatzek, S., Niehaus, K., Boller, T., and Felix, G. (2004). The N terminus of bacterial elongation factor Tu elicits innate immunity in Arabidopsis plants. Plant Cell 16, 3496–3507.

24. Xu, F., and Yu, F. (2023). Sensing and regulation of plant extracellular pH. Trends Plant Sci. 28,1422–1437

25. Li, L., Verstraeten, I., Roosjen, M., Takahashi, K., Rodriguez, L., Merrin, J., Chen, J., Shabala, L., Smet, W., Ren, H., et al. (2021). Cell surface and intracellular auxin signalling for H^+^ fluxes in root growth. Nature 599, 273–277.

26. Lin, W., Zhou, X., Tang, W., Takahashi, K., Pan, X., Dai, J., Ren, H., Zhu, X., Pan, S., Zheng, H., et al. (2021). TMK-based cell-surface auxin signalling activates cell-wall acidification. Nature 599, 278–282.

27. Caesar, K., Elgass, K., Chen, Z., Huppenberger, P., Witthoft, J., Schleifenbaum, F., Blatt, M.R., Oecking, C., and Harter, K. (2011). A fast brassinolide-regulated response pathway in the plasma membrane of *Arabidopsis thaliana*. Plant J. 66, 528–540.

28. Fuglsang, A.T., Kristensen, A., Cuin, T.A., Schulze, W.X., Persson, J., Thuesen, K.H., Ytting, C.K., Oehlenschlaeger, C.B., Mahmood, K., Sondergaard, T.E., et al. (2014). Receptor kinase-mediated control of primary active proton pumping at the plasma membrane. Plant J. 80, 951–964.

29. Haruta, M., Sabat, G., Stecker, K., Minkoff, B.B., and Sussman, M.R. (2014). A peptide hormone and its receptor protein kinase regulate plant cell expansion. Science 343, 408–411.

30. Elmore, J.M., and Coaker, G. (2011). The role of the plasma membrane H^+^-ATPase in plant-microbe interactions. Mol. Plant 4, 416–427.

31. Falhof, J., Pedersen, J.T., Fuglsang, A.T., and Palmgren, M. (2016). Plasma membrane H^+^-ATPase regulation in the center of plant physiology. Mol. Plant 9, 323–337.

32. Diaz-Ardila, H.N., Gujas, B., Wang, Q., Moret, B., and Hardtke, C.S. (2023). pH-dependent CLE peptide perception permits phloem differentiation in Arabidopsis roots. Curr. Biol. 33 597–605.e3.

33. Liu, L., Song, W., Huang, S., Jiang, K., Moriwaki, Y., Wang, Y., Men, Y., Zhang, D., Wen, X., Han, Z., et al. (2022). Extracellular pH sensing by plant cell-surface peptide-receptor complexes. Cell 185, 3341–3355.

34. Zhai, K., Rhodes, J., and Zipfel, C. (2024). A peptide-receptor module links cell wall integrity sensing to pattern-triggered immunity. Nat. Plants 10, 2027–2037.

35. Han, M., Yang, H., Yu, G., Jiang, P., You, S., Zhang, L., Lin, H., Liu, J., and Shu, Y. (2022). Application of non-invasive micro-test technology (NMT) in environmental fields: a comprehensive review. Ecotox Environ. Safe 240, 113706.

36. Elke Barbez, K.D., Angelika Gaidora, Thomas Lendl, and Wolfgang Busch (2017). Auxin steers root cell expansion via apoplastic pH regulation in *Arabidopsis thaliana*. Proc. Natl. Acad. Sci. U.S.A. 114, E4884–E4893.

37. Tang, J., Han, Z., Sun, Y., Zhang, H., Gong, X., and Chai, J. (2015). Structural basis for recognition of an endogenous peptide by the plant receptor kinase PEPR1. Cell Res. 25, 110–120.

38. Bohm, H., Albert, I., Oome, S., Raaymakers, T.M., Van den Ackerveken, G., and Nurnberger, T. (2014). A conserved peptide pattern from a widespread microbial virulence factor triggers pattern-induced immunity in Arabidopsis. PLoS Pathog. 10, e1004491.

39. Perraki, A., DeFalco, T.A., Derbyshire, P., Avila, J., Sere, D., Sklenar, J., Qi, X., Stransfeld, L., Schwessinger, B., Kadota, Y., et al. (2018). Phosphocode-dependent functional dichotomy of a common co-receptor in plant signalling. Nature 561, 248–252.

40. Ueno, K., Kinoshita, T., Inoue, S., Emi, T., and Shimazaki, K. (2005). Biochemical characterization of plasma membrane H^+^-ATPase activation in guard cell protoplasts of *Arabidopsis thaliana* in response to blue light. Plant Cell Physiol. 46, 955–963.

41. Beck, M., Wyrsch, I., Strutt, J., Wimalasekera, R., Webb, A., Boller, T., and Robatzek, S. (2014). Expression patterns of flagellin sensing 2 map to bacterial entry sites in plant shoots and roots. J. Exp. Botany 65, 6487–6498.

42. Haruta, M., Burch, H.L., Nelson, R.B., Barrett-Wilt, G., Kline, K.G., Mohsin, S.B., Young, J.C., Otegui, M.S., and Sussman, M.R. (2010). Molecular characterization of mutant Arabidopsis plants with reduced plasma membrane proton pump activity. J. Biol. Chem. 285, 17918–17929.

43. Merlot, S., Leonhardt, N., Fenzi, F., Valon, C., Costa, M., Piette, L., Vavasseur, A., Genty, B., Boivin, K., Muller, A., et al. (2007). Constitutive activation of a plasma membrane H^+^-ATPase prevents abscisic acid-mediated stomatal closure. EMBO J. 26, 3216–3226.

44. Takahashi, K., Hayashi, K., and Kinoshita, T. (2012). Auxin activates the plasma membrane H^+^-ATPase by phosphorylation during hypocotyl elongation in Arabidopsis. Plant Physiol. 159, 632–641.

45. Pei, D., Hua, D., Deng, J., Wang, Z., Song, C., Wang, Y., Wang, Y., Qi, J., Kollist, H., Yang, S., et al. (2022). Phosphorylation of the plasma membrane H^+^-ATPase AHA2 by BAK1 is required for ABA-induced stomatal closure in Arabidopsis. Plant Cell 34, 2708–2729.

46. Snoeck, S., Lee, H.K., Schmid, M.W., Bender, K.W., Neeracher, M.J., Fernandez-Fernandez, A.D., Santiago, J., and Zipfel, C. (2024). Leveraging coevolutionary insights and AI-based structural modeling to unravel receptor-peptide ligand-binding mechanisms. Proc. Natl. Acad. Sci. U. S. A. 121, e2400862121.

47. Jia, F., Xiao, Y., Feng, Y., Yan, J., Fan, M., Sun, Y., Huang, S., Li, W., Zhao, T., Han, Z., et al. (2024). N-glycosylation facilitates the activation of a plant cell-surface receptor. Nat. Plants 10, 2014–2026.

48. Wu, H., Wan, L., Liu, Z., Jian, Y., Zhang, C., Mao, X., Wang, Z., Wang, Q., Hu, Y., Xiong, L., et al. (2024). Mechanistic study of SCOOPs recognition by MIK2-BAK1 complex reveals the role of N-glycans in plant ligand-receptor-coreceptor complex formation. Nat. Plants 10, 1984–1998.

49. Zhang, J., Li, W., Xiang, T., Liu, Z., Laluk, K., Ding, X., Zou, Y., Gao, M., Zhang, X., Chen, S., et al. (2010). Receptor-like cytoplasmic kinases integrate signaling from multiple plant immune receptors and are targeted by a *Pseudomonas syringae* effector. Cell Host Microbe 7, 290–301.

50. Lu, D., Wu, S., Gao, X., Zhang, Y., Shan, L., and He, P. (2010). A receptor-like cytoplasmic kinase, BIK1, associates with a flagellin receptor complex to initiate plant innate immunity. Proc. Natl. Acad. Sci. U. S. A. 107, 496-501.

51. Wenwei Lin, B.L., Dongping Lu, Sixue Chen, Ning Zhu, Ping He, and Libo Shan (2014). Tyrosine phosphorylation of protein kinase complex BAK1/BIK1 mediates Arabidopsis innate immunity. Proc. Natl. Acad. Sci. U. S. A. 111, 3632–3637.

52. Pallucca, R., Visconti, S., Camoni, L., Cesareni, G., Melino, S., Panni, S., Torreri, P., and Aducci, P. (2014). Specificity of epsilon and non-epsilon isoforms of arabidopsis 14-3-3 proteins towards the H^+^-ATPase and other targets. PloS One 9, e90764.

53. Thor, K., Jiang, S., Michard, E., George, J., Scherzer, S., Huang, S., Dindas, J., Derbyshire, P., Leitao, N., DeFalco, T.A., et al. (2020). The calcium-permeable channel OSCA1.3 regulates plant stomatal immunity. Nature 585, 569–573.

54. Li, L., Li, M., Yu, L., Zhou, Z., Liang, X., Liu, Z., Cai, G., Gao, L., Zhang, X., Wang, Y., et al. (2014). The FLS2-associated kinase BIK1 directly phosphorylates the NADPH oxidase RbohD to control plant immunity. Cell Host Microbe 15, 329–338.

55. Liang, X., Ding, P., Lian, K., Wang, J., Ma, M., Li, L., Li, L., Li, M., Zhang, X., Chen, S., et al. (2016). Arabidopsis heterotrimeric G proteins regulate immunity by directly coupling to the FLS2 receptor. Elife 5, e13568.

56. Wang, W., Fei, Y., Wang, Y., Song, B., Li, L., Zhang, W., Cheng, H., Zhang, X., Chen, S., and Zhou, J.M. (2023). SHOU4/4L link cell wall cellulose synthesis to pattern-triggered immunity. New Phytol. 238 (4), 1620–1635.

57. Kadota, Y., Sklenar, J., Derbyshire, P., Stransfeld, L., Asai, S., Ntoukakis, V., Jones, J.D., Shirasu, K., Menke, F., Jones, A., et al. (2014). Direct regulation of the NADPH oxidase RBOHD by the PRR-associated kinase BIK1 during plant immunity. Mol. Cell 54, 43–55.

58. Dindas, J., DeFalco, T.A., Yu, G., Zhang, L., David, P., Bjornson, M., Thibaud, M.C., Custodio, V., Castrillo, G., Nussaume, L., et al. (2022). Direct inhibition of phosphate transport by immune signaling in Arabidopsis. Curr. Biol. 32, 488–495 e485.

59. Chu, J., Monte, I., DeFalco, T.A., Koster, P., Derbyshire, P., Menke, F.L.H., and Zipfel, C. (2023). Conservation of the PBL-RBOH immune module in land plants. Curr. Biol. 33, 1130–1137 e1135.

60. DeFalco, T.A., Anne, P., James, S.R., Willoughby, A.C., Schwanke, F., Johanndrees, O., Genolet, Y., Derbyshire, P., Wang, Q., Rana, S., et al. (2022). A conserved module regulates receptor kinase signalling in immunity and development. Nat. Plants 8, 356–365.

61. Tian, W., Hou, C., Ren, Z., Wang, C., Zhao, F., Dahlbeck, D., Hu, S., Zhang, L., Niu, Q., Li, L., et al. (2019). A calmodulin-gated calcium channel links pathogen patterns to plant immunity. Nature 572, 131–135.

62. Kong, L., Ma, X., Zhang, C., Kim, S.I., Li, B., Xie, Y., Yeo, I.C., Thapa, H., Chen, S., Devarenne, T.P., et al. (2024). Dual phosphorylation of DGK5-mediated PA burst regulates ROS in plant immunity. Cell 187 (3), 609–623. e21.

63. Qi, F., Li, J., Ai, Y., Shangguan, K., Li, P., Lin, F., and Liang, Y. (2024). DGK5β-derived phosphatidic acid regulates ROS production in plant immunity by stabilizing NADPH oxidase. Cell Host Microbe 32, 425–440 e427.

64. Liang, X., Ma, M., Zhou, Z., Wang, J., Yang, X., Rao, S., Bi, G., Li, L., Zhang, X., Chai, J., et al. (2018). Ligand-triggered de-repression of Arabidopsis heterotrimeric G proteins coupled to immune receptor kinases. Cell Res. 28, 529–543.

65. Wortmann, A., He, Y., Christensen, M.E., Linn, M., Lumley, J.W., Pollock, P.M., Waterhouse, N.J., and Hooper, J.D. (2011). Cellular settings mediating Src Substrate switching between focal adhesion kinase tyrosine 861 and CUB-domain-containing protein 1 (CDCP1) tyrosine 734. J. Biol. Chem. 286, 42303–42315.

66. Clark, L.K., and Cullati, S.N. (2025). Activation is only the beginning: mechanisms that tune kinase substrate specificity. Biochem. Soc. Trans. 53 (01), 145–159.

67. Watkins, J.M., Montes, C., Clark, N.M., Song, G., Oliveira, C.C., Mishra, B., Brachova, L., Seifert, C.M., Mitchell, M.S., Yang, J., et al. (2024). Phosphorylation dynamics in a flg22-Induced, G protein-dependent network reveals the AtRGS1 phosphatase. Mol. Cell Proteom. 23, 100705.

68. Liu, Z., Jia, Y., Ding, Y., Shi, Y., Li, Z., Guo, Y., Gong, Z., and Yang, S. (2017). Plasma membrane CRPK1-mediated phosphorylation of 14-3-3 proteins induces their nuclear import to fine-tune CBF signaling during cold response. Mol. Cell 66, 117–128 e115.

69. Salvi, M., Trashi, E., Cozza, G., Franchin, C., Arrigoni, G., and Pinna, L.A. (2012). Investigation on PLK2 and PLK3 substrate recognition. Biochim. Biophys. Acta 1824, 1366–1373.

70. Lal, N.K., Nagalakshmi, U., Hurlburt, N.K., Flores, R., Bak, A., Sone, P., Ma, X., Song, G., Walley, J., Shan, L., et al. (2018). The receptor-like cytoplasmic kinase BIK1 localizes to the nucleus and regulates defense hormone expression during plant innate immunity. Cell Host Microbe 23, 485–497 e485.

71. Kim, S., Park, J., Jeon, B.W., Hwang, G., Kang, N.Y., We, Y., Park, W.Y., Oh, E., and Kim, J. (2021). Chemical control of receptor kinase signaling by rapamycin-induced dimerization. Mol. Plant 14, 1379–1390.

72. Mühlenbeck, H., Tsutsui, Y., Lemmon, M. A., Bender, K. W., & Zipfel, C. (2024). Allosteric activation of the co-receptor BAK1 by the EFR receptor kinase initiates immune signaling. eLife, 12, RP92110.

73. Stegmann, M., Monaghan, J., Smakowska-Luzan, E., Rovenich, H., Lehner, A., Holton, N., Belkhadir, Y., and Zipfel, C. The receptor kinase FER is a RALF-regulated scaffold controlling plant immune signaling. Science, 355, 287–289.

74. Tateno, M., Brabham, C., and DeBolt, S. (2016). Cellulose biosynthesis inhibitors - a multifunctional toolbox. J. Exp. Bot. 67, 533–542.

75. Engelsdorf, T., Gigli-Bisceglia, N., Veerabagu, M., McKenna, J.F., Vaahtera, L., Augstein, F., Van der Does, D., Zipfel, C., and Hamann, T. (2018). The plant cell wall integrity maintenance and immune signaling systems cooperate to control stress responses in *Arabidopsis thaliana*. Sci. Signal. 11, eaao3070.

76. Denison, F.C., Paul, A.L., Zupanska, A.K., and Ferl, R.J. (2011). 14-3-3 proteins in plant physiology. Semin Cell Dev Biol. 22, 720–727.

77. Wolf, S. (2022). Cell wall signaling in plant development and defense. Annu. Rev. Plant Biol. 73, 323–353.

78. Kesten, C., Gamez-Arjona, F.M., Menna, A., Scholl, S., Dora, S., Huerta, A.I., Huang, H.Y., Tintor, N., Kinoshita, T., Rep, M., et al. (2019). Pathogen-induced pH changes regulate the growth-defense balance in plants. EMBO J. 38, e101822.

79. Yu, K., Liu, Y., Tichelaar, R., Savant, N., Lagendijk, E., van Kuijk, S.J.L., Stringlis, I.A., van Dijken, A.J.H., Pieterse, C.M.J., Bakker, P., et al. (2019). Rhizosphere-associated pseudomonas suppress local root immune responses by gluconic acid-mediated lowering of environmental pH. Curr. Biol. 29, 3913–3920 e3914.

80. Geilfus, C.M. (2017). The pH of the apoplast: dynamic factor with functional impact under stress. Mol. Plant 10, 1371–1386.

81. Tsai, H.H., and Schmidt, W. (2021). The enigma of environmental pH sensing in plants. Nat. Plants 7, 106–115.

82. Leicher, H., Schade, S., Huebbers, J.W., Munzert-Eberlein, K.S., Haljiti, G., Ludwig, C., Mueller, M., Kinoshita, T., Engelsdorf, T., Hueckelhoven, R., et al. (2025). Endogenous RALF peptide function is required for powdery mildew host colonization. Preprint at bioRxiv 10.1101/2025.01.30.635691.

83. Fuglsang, A.T., Guo, Y., Cuin, T.A., Qiu, Q., Song, C., Kristiansen, K.A., Bych, K., Schulz, A., Shabala, S., Schumaker, K.S., et al. (2007). Arabidopsis protein kinase PKS5 inhibits the plasma membrane H^+^-ATPase by preventing interaction with 14-3-3 protein. Plant Cell 19, 1617–1634.

84. Nühse, T.S., Bottrill, A.R., Jones, A.M., and Peck, S.C. (2007). Quantitative phosphoproteomic analysis of plasma membrane proteins reveals regulatory mechanisms of plant innate immune responses. Plant J. 51, 931–940.

85. Sedlov, I.A., and Sluchanko, N.N. (2025). The big, mysterious world of plant 14-3-3 proteins. Biochemistry (Mosc) 90, S1–S35.

86. Van Der Hoeven, P.C., Van Der Wal, J.C., Ruurs, P., Van Dijk, M.C., and Van Blitterswijk, J. (2000). 14-3-3 isotypes facilitate coupling of protein kinase C-zeta to Raf-1: negative regulation by 14-3-3 phosphorylation. Biochem. J. 345, 297–306.

87. Powell, D.W., Rane, M.J., Joughin, B.A., Kalmukova, R., Hong, J.H., Tidor, B., Dean, W.L., Pierce, W.M., Klein, J.B., Yaffe, M.B., et al. (2003). Proteomic identification of 14-3-3zeta as a mitogen-activated protein kinase-activated protein kinase 2 substrate: role in dimer formation and ligand binding. Mol. Cell Biol. 23, 5376–5387.

88. Dong, X., Feng, F., Li, Y., Li, L., Chen, S., and Zhou, J.M. (2023). 14-3-3 proteins facilitate the activation of MAP kinase cascades by upstream immunity-related kinases. Plant Cell. 35, 2413–2428.

89. Roux, M., Schwessinger, B., Albrecht, C., Chinchilla, D., Jones, A., Holton, N., Malinovsky, F.G., Tor, M., de Vries, S., and Zipfel, C. (2011). The Arabidopsis leucine-rich repeat receptor-like kinases BAK1/SERK3 and BKK1/SERK4 are required for innate immunity to hemibiotrophic and biotrophic pathogens. Plant Cell 23, 2440–2455.

90. Hayashi, Y., Nakamura, S., Takemiya, A., Takahashi, Y., Shimazaki, K., and Kinoshita, T. (2010). Biochemical characterization of in vitro phosphorylation and dephosphorylation of the plasma membrane H^+^-ATPase. Plant Cell Physiol. 51, 1186–1196.

91. Platre, M.P., Noack, L.C., Doumane, M., Bayle, V., Simon, M.L.A., Maneta-Peyret, L., Fouillen, L., Stanislas, T., Armengot, L., Pejchar, P., et al. (2018). A combinatorial lipid code shapes the electrostatic landscape of plant endomembranes. Dev. Cell 45, 465–480 e411.

92. Perez-Riverol, Y., Csordas, A., Bai, J., Bernal-Llinares, M., Hewapathirana, S., Kundu, D.J., Inuganti, A., Griss, J., Mayer, G., Eisenacher, M., et al. (2019). The PRIDE database and related tools and resources in 2019: improving support for quantification data. Nucleic Acids Res. 47, D442–D450.

93. Chiasson, D., Gimenez-Oya, V., Bircheneder, M., Bachmaier, S., Studtrucker, T., Ryan, J., Sollweck, K., Leonhardt, H., Boshart, M., Dietrich, P., et al. (2019). A unified multi-kingdom Golden Gate cloning platform. Sci. Rep. 9, 10131.

94. Zhang, X., Henriques, R., Lin, S.S., Niu, Q.W., and Chua, N.H. (2006). Agrobacterium-mediated transformation of *Arabidopsis thaliana* using the floral dip method. Nat. Protoc. 1, 641–646.

95. Yoo, S.-D., Cho, Y.-H., and Sheen, J. (2007). Arabidopsis mesophyll protoplasts: a versatile cell system for transient gene expression analysis. Nat. Protoc. 2, 1565–1572.

96. Iyer, K., Bürkle, L., Auerbach, D., Thaminy, S., Dinkel, M., Engels, K., and Stagljar, I. (2005). Utilizing the split-ubiquitin membrane yeast two-hybrid system to identify protein-protein interactions of integral membrane proteins. Sci. STKE 2005, pl3.

97. Manhães, A.M.E.d.A., de Oliveira, M.V.V., and Shan, L. (2015). Establishment of an efficient virus-induced gene silencing (VIGS) assay in Arabidopsis by agrobacterium-mediated rubbing infection. In plant gene silencing: methods and protocols, K.S. Mysore and M. Senthil-Kumar, eds. (New York, NY: Springer New York), pp. 235-241.

98. Shabala, S.N., Newman, I.A., and Morris, J. (1997) Oscillations in H^+^ and Ca^2+^ ion fluxes around the elongation region of corn roots and effects of external pH. Plant Physiol. 113, 111–118.

